# Variability by region and method in human brain sodium concentrations estimated by ^23^Na magnetic resonance imaging: a meta-analysis

**DOI:** 10.1101/2022.11.02.514873

**Authors:** Ben Ridley, Filomena Morsillo, Wafaa Zaaraoui, Francesco Nonino

## Abstract

Sodium imaging (^23^Na-MRI) is of interest in neurological conditions given potential sensitivity to the physiological and metabolic status of tissues. Benchmarks have so far been restricted to parenchyma or grey/white matter (GM/WM). We investigate (1) the availability of evidence, (2) regional pooled estimates and (3) variability attributable to regional/methodology.

MEDLINE literature search for Tissue sodium concentration (TSC) measured in specified ‘healthy’ brain regions returned 127 reports plus 278 retrieved from bibliographies. 28 studies met inclusion criteria, including 400 individuals. Reporting variability led to nested data structure, so we used multilevel meta-analysis and a random effects model to pool effect sizes.

The pooled mean from 141 TSC estimates was 40.51 mM (95% CI: 37.59 - 43.44; p< 0.001, I^2^_Total=_99.4%). Tissue as a moderator was significant (F^2^_14_=65.34, p-val < .01). Six sub-regional pooled means with requisite statistical power were derived. We were unable to consider most methodological and demographic factors sought because of non-reporting, but each factor included beyond tissue improved model fit. Significant residual heterogeneity remained.

The current estimates provide an empirical point of departure for better understanding in ^23^Na-MRI. Improving on current estimates supports: (1) larger, more representative data collection/sharing, including (2) regional data, and (3) agreement on full reporting standards.

## Introduction

Sodium magnetic resonance imaging (^23^Na-MRI) is of interest as a ‘quantitative’ imaging modality. Using a reference of known concentration **(Figure 1a)**, measured signal (M0) can be converted from arbitrary signal intensity to a quantitative scale (millimolars, mM). As a candidate for metabolic imaging in particular, the advantages of ^23^Na-MRI include: 1) it natively produces 3D, whole-brain, voxel-based data and is not restricted to pre-defined volume-of-interest analyses, and 2) the fact it requires no contrast agents, meaning contraindications are the same as for conventional proton (^1^H) MRI. Ionic homeostasis is a pre-requisite for proper cellular functioning, with sodium in the nervous system being critical in trans-membrane transport, osmotic and electrostatic regulation and the generation/propagation of action potentials^1–3^. As such, the non-invasive, in vivo measurement of sodium concentration by ^23^Na-MRI is of interest in the context of neuro-oncological^4–6^, neurodegenerative^7–9^, demyelinating^10–19^ and cerebrovascular^20^ conditions, and in both physiological and pathological neuronal activity^21–25^.

**Figure 1.**
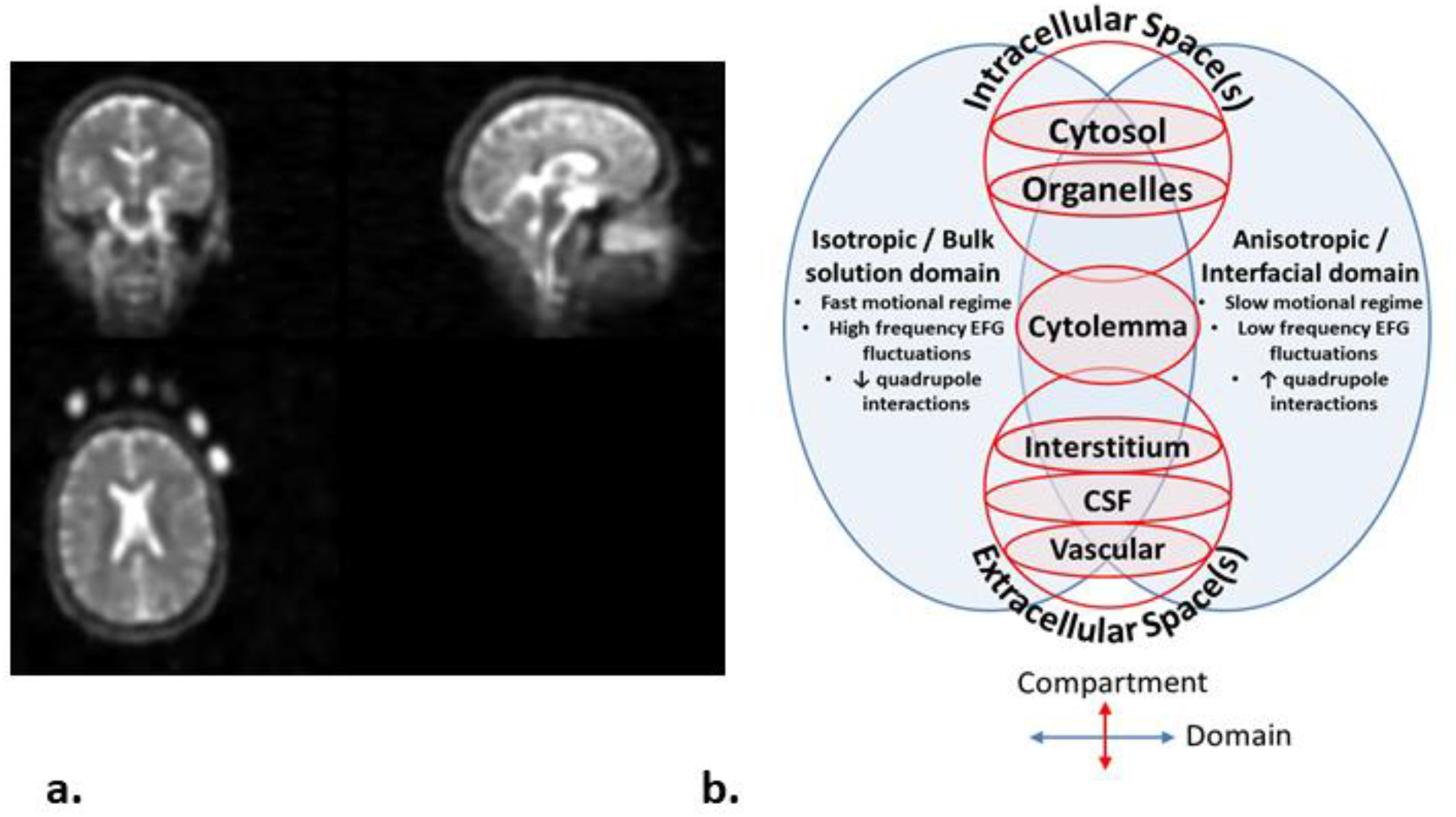
^23^Na magnetic resonance imaging. a) An exemplar 23Na-MRI brain image, with external calibration phantoms of differing concentrations visible on the bottom left axial image. Calibration can also be done relative to internal references such as vitreous humor or cerebrospinal fluid in the ventricles. b) Schematic Venn diagram (following the nomenclature in Springer^28^) showing physical domains (Blue) where nuclei share similar nuclear magnetic resonance properties, and biological compartments (Red) found in brain tissue. These are arrayed along two axes to indicate their orthogonality: neither physical domain is unique to any given biological compartment. ^23^Na-MRI TSC estimates are a weighted average influenced by concentrations, volumes and microstructure in a range of environments including the intracellular (neuronal and glial cytosol and organelles), extracellular (interstitial and vascular spaces.) and membranous (cyto/axolemmal) contributions ^28^. The vast majority of *in vivo* sodium is present in metal-aquo complexes with a tetrahedral hydration shell surrounding the ^23^Na ion, with a much smaller population bound to macromolecular loci^29^. The domain on the left corresponds to the situation in bulk solution, where magnetic and electric fields average out to become isotropic. An anisotropic domain (right) pertains at the interface of/with macromolecules and/or lipid assemblies, where the surface experienced by a diffusing ion is not randomly orientated and the resulting electric field gradient (EFG) fluctuations do not average to zero. ^23^Na is a quadropolar ion (spin=3/2) that, under the influence of a magnetic field, exhibits four energy levels with three possible single quantum transitions, one central and two satellites each contributing to relaxation^30^. In the context of isotropic domains, *i*.*e*. aqueous environments with rapid motions, quadrupole interactions are minimal and all transitions occur approximately at the same decay time resulting in a MRI-visible monoexponential decay curve ^30^. In anisotropic domains, where motions are slowed, the non-spherical distribution of the electric charge of the sodium nucleus permits interaction with anisotropic electric fields of the charged groups on the macromolecular anions. Thus, quadrupole interactions are non-zero and biexponential relaxation is observed ^29,30,32,33^ with the satellite transitions showing faster decay than the central transition.

In practice, assigning a single imaging parameter to a voxel in an MRI image of biological tissue is an oversimplification. This is the case both for the weighted average referred to as ‘total’ or ‘tissue sodium concentration’ (TSC) in ^23^Na-MRI, as well as ‘conventional’ MRI contrasts targeting tissue water protons (^1^H) such as diffusion or T_2_ measurements^26,27^. Image sampling/tissue fraction effects are one reason, where diverse tissue types such as white and grey matter (WM and GM) and cerebrospinal fluid (CSF) contribute to measured signal within a single voxel. Even within a given tissue type MRI cannot resolve the sub-cellular compartments/organelles that, in the case of ^23^Na-MRI, can be said to actually have a specific concentration^28^. The measurement of a given voxel will include, at a minimum, intracellular and extracellular compartments with their own concentrations, volumes and microstructure (**Figure 1b**).

Physical behaviour of sodium atoms in biological media and the complexities of measuring them with MRI present other challenges. Relative to tissue water protons, the lower MR sensitivity and abundance of ^23^Na result in lower signal to noise ratios and larger voxel sizes, exacerbating tissue fraction effects^34,35. 23^Na-MRI pulse sequences with ultra-fast echo times (TE) and non-cartesian sampling schemes that compensate for the short, biexponential transverse relaxation of ^23^Na nuclei ^34–37^ have broader point spread functions (PSF) and inter-voxel spill-over is a greater issue. These partial volume effect (PVE) issues are the target of growing attempts to develop or import PVE correction techniques, such as those adapted from positron emission tomography imaging (PET) ^12,34–37^. Correction techniques beg the question of benchmarks for correction algorithms to target.

A range of tissue volume models to understand and validate ^23^Na-MRI-derived concentrations have been proposed. The ‘canonical’ model describes two compartments: a large volume/low concentration (variously 10-15mM^5,6,10,20,38,39^) intracellular space and an extracellular space of smaller volume but higher concentration (140mM^5, 6,10,20^ or 145^38,39^ mM). These figures lead to a general estimate for overall brain tissue (parenchyma) of about 37-45mM^6, 12,20^. Model-based estimates beyond a general figure for parenchyma or gross tissue divisions like GM or WM are largely lacking^40^, particularly because the cellular data required to elaborate beyond this are for the most part not available. More broadly, histology is an invaluable tool but should not be naively taken as the absolute gold-standard for MR features both because it is not necessarily sensitive to the same properties as MR^27^ and because various techniques suffer from their own limitations in regarding compartmental concentrations, ecological validity given preparation effects, and limited spatial, temporal and cross-species sampling^28^.

The positioning of ^23^Na-MRI as a putative ‘quantitative’ method, implies that concentration measures should converge toward a ‘true’ estimate for a given sample, modulo statistical and methodological effects. If ^23^Na-MRI is sensitive to the physiological state of tissues – a key assumption motivating its use in the context of neurological conditions – regional variability in measured TSC concentrations should also be expected. Conversely, positing a single parenchymal concentration as sufficient to characterise all regions implies a limit on detectable differences between individuals with and without neurological conditions, given known variation in factors like fluid fractions^41,42^, distributions of cellular types and architectures^43,44^ and macromolecular content^45–47^.

In this context, meta-analytic approaches are another means to synthesize evidence and identify impediments and progress towards consensus. Meta-analysis aims to estimate the true effect size (including central tendency measures like the mean) based on the combination of observed effect sizes taken from several empirical samples, while trying to account for sample and study variability^48^. As such, we sought to apply meta-analytic measures to investigate the existing literature on estimates of TSC in human brain regions. We aimed to address (1) the range of available evidence in the form of regional TSC estimates in the literature, (2) the possibility of consensus estimates of concentration in various brain regions, and (3) the extent to which methodological and anatomical factors contribute to variation in measured TSC.

## Results

### Search Results

A search (see Methods) of MEDLINE dated 12/7/2021 returned 127 records, and we identified an additional 278 records by examining the bibliographies of recovered records and the ‘cited by’ function on the PubMed website. These records underwent screening of titles and abstracts, and the remaining texts underwent full text assessment for inclusion (see Preferred Reporting Items for Systematic Reviews and Meta-Analyses (PRISMA) flow diagram **Figure 2**, and **Supplementary Tables 1-2** for PRISMA checklists). From an overall total of 405 records, 44 records were screened from further consideration based on title/abstract. 361 records were sought for full text retrieval, of which 329 were excluded with the main reasons being a focus on non-brain tissue or the lack of a sodium measurement in the form of a concentration estimate. We were unable to access four records. See **Supplementary Table 3** for a full list of records identified and reasons for exclusion. Inclusion criteria were met by 28 reports.

**Figure 2.**
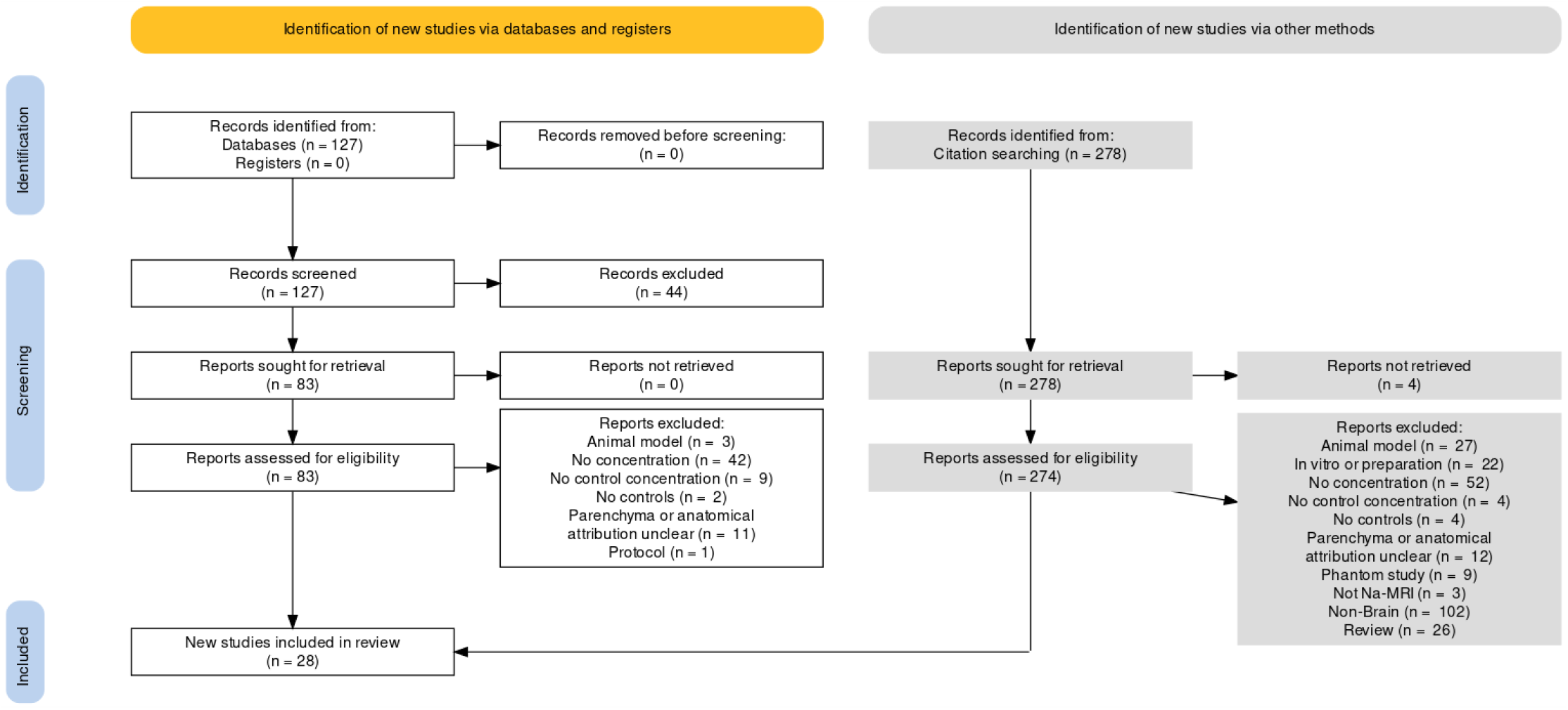
PRISMA flow diagram for search performed 12/7/2021. Generated with the PRISMA 2020 app^49^.

### Included studies

We included 28 reports containing measurements of total sodium concentration in healthy controls of specified human brain regions or tissue divisions (other than ‘parenchyma’) (**Table 1**). Nominally, 400 healthy controls in total were included in these reports, the mean number per report being 14.3 individuals (range: 4-45, S.D: 11.1). All but one included report was published in the last ten years, and all but three since 2010 (**Figure 3a**).

**Table 1.** Characteristics of included studies.

| Paper | Ref | Tesla | Sequence | Calibration<sup>a</sup> | TR<sup>b</sup><br>(ms) | TE<sup>c</sup><br>(ms) | Voxel<br>volume<br>(mm<sup>3</sup>)<sup>d</sup> | N | N.<br>Female | Age<br>(years)<sup>e</sup> | Comparison<br>group |
| --- | --- | --- | --- | --- | --- | --- | --- | --- | --- | --- | --- |
| Winkler et al 1989 | <sup>51</sup> | 1.5 | GRE | Int. VH | 98 | 3 | 490 | 4 | NR | 20-35 | None |
| Ouwerkerk et al 2003 | <sup>5</sup> | 1.5 | TPI | External | 120 | 0,37 | 39.3 | 9 | 3 | 22-63 | Tumour |
| Thulborn et al 2005 | <sup>52</sup> | 3 | TPI | External | 100 | 0,3 | 125 | 5 | NR | NR | Stroke |
| Inglese et al 2010 | <sup>10</sup> | 3 | Radial | External | 120 | 0,05 | 64 | 13 | 10 | 36.7,<br>26–60 | MS |
| Lu et al 2010 | <sup>38</sup> | 3 | FlexTPI | External | 160 | 0,36 | 125 | 5 | 1 | 32.4 ±<br>8.9 | None |
| Reetz et al 2012 | <sup>8</sup> | 4 | SPRITE | External | 10 | 0,3 | 64 | 13 | 6 | 44.9 ±<br>9.9 | HD |
| Qian et al 2012 | <sup>53</sup> | 7 | AWSOS | Int. CSF | 100 | 0,5 | 2.9 | 5 | 5 | 20–48 | None |
| Zaaraoui et 2012 | <sup>11</sup> | 3 | DA Radial | External | 120 | 0,2 | 46.6 | 15 | 12 | 30,<br>20–54 | MS |
| Paling et al 2013 | <sup>12</sup> | 3 | Radial | External | 120 | 0,27 | 64 | 27 | 16 | 42.9<br>±11.3 | MS |
| Maarouf et al 2014 | <sup>13</sup> | 3 | DA Radial | External | 120 | 0,2 | 46.6 | 15 | NR | 30,<br>21-54 | MS |
| Mirkes et al 2015 | <sup>54</sup> | 9.4 | AWSOS | External | 150 | 0,3 | 5 | 5 | 1 | 29±4 | None |
| Niesporak et al 2015 | <sup>55</sup> | 7 | DA Radial | External | 150 | 0,45 | 27 | 4 | 1 | 26 ± 2 | None |
| Eisele et al 2016 | <sup>15</sup> | 3 | DA Radial | External | 60 | 0,22 | 46.6 | 10 | 5 | 33,<br>23–53 | MS |
| Petracca et al 2016 | <sup>16</sup> | 7 | GRE | External | 150 | 6,8 | 125 | 17 | 8 | 46.16 ±<br>11.65 | MS |
| Thulborn et al 2016 | <sup>39</sup> | 9.4 | FlexTPI | External | 160 | 0,26 | 42.9 | 45 | NR | 48 ± 19 | None |
| Maarouf et al 2017 | <sup>14</sup> | 3 | DA Radial | External | 120 | 0,2 | 46.7 | 31 | 15 | 35.7<br>± 12.4 | MS |
| Eisele et al 2017 | <sup>17</sup> | 3 | DA Radial | External | 60 | 0,22 | 46.7 | 6 | 5 | 42 ± 10 | MS |
| Ridley et al 2018 | <sup>56</sup> | 7 | DA Radial | External | 120 | 0,3 | 42.9 | 13 | 5 | 23.9 ±<br>3.6 | None |
| Worthoff et al 2018 | <sup>57</sup> | 4 | SISTINA | Int. VH | 150 | 0,36 | 216 | 40 | 16 | 19–70 | None |
| Reimer et al 2019 | <sup>58</sup> | 3 | Cones | External | 100 | 0,5 | 64 | 11 | 3 | 32 ± 6 | None |
| Driver et al 2019 | <sup>59</sup> | 4.7 | TPI | Int. VH | 85 | 0,11 | 65.5 | 9 | 5 | 30 ± 6 | None |
| Liao et al 2019 | <sup>60</sup> | 3 | TPI | Int. CSF | 160 | 0,4 | 40.7 | 8 | 3 | 25-32 | None |
| Meyer et al 2019a | <sup>61</sup> | 3 | DA Radial | External | 120 | 0,2 | 46.7 | 12 | 8 | 31±8.3 | None |
| Meyer et al 2019b | <sup>24</sup> | 3 | DA Radial | External | 120 | 0,2 | 64 | 12 | 12 | 34.3 ± 10.7 | Migraine |
| Kim et al 2020 | <sup>37</sup> | 7 | GRE | Int. CSF | 100 | 4 | 64 | 8 | 0 | 20-35 | None |
| Gerhalter et al 2021 | <sup>62</sup> | 3 | FLORET | Int. VH | 100 | 0,2 | 216 | 19 | 12 | 31,4 ± 7,5 | TBI |
| Brownlee et al 2019 | <sup>19</sup> | 3 | Cones | External | 120 | 0,22 | 27 | 34 | 23 | 35,5 ± 10,1 | MS |
| Schneider et al 2021 | <sup>63</sup> | 7 | DA Radial | Int. CSF | 100 | 0,35 | 8 | 5 | 3 | 28,4 ± 6,5 | None |

a. Int. – internal, Ext. – External, CSF – cerebrospinal fluid, VH – vitreous humour (eyes)
b. Repetition time
c. Echo time, where multiple the shortest TE was used in analysis
d. x,y,z product of nominal resolution, see **Supplementary Table 7** for specific values.
e. NR =not reported, other values are presented as per original reports in ranges (XX-XX), mean\_± standard deviation, except for Zaaraoui et 2012 and Maarouf et al 2014 which report median and range and Inglese et al 2010 and Eisele et al 2016 which report mean and range.

**Figure 3:**
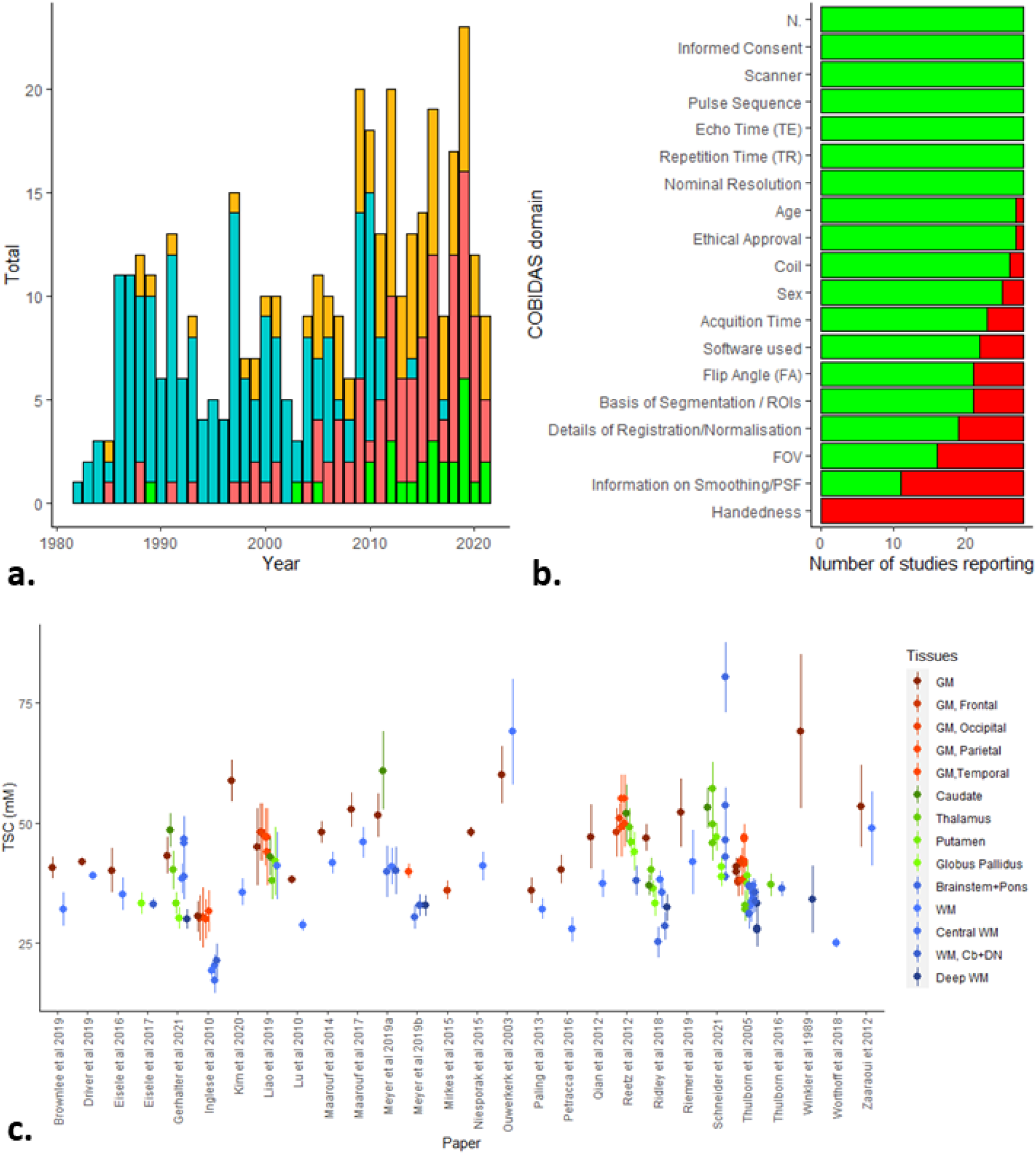
Descriptive plots for identified and included studies. **(A)** Literature search results 1980-2021 by year of publication. Chart includes total papers published for a given year (orange), publications identified by searching bibliographies (cyan), papers identified through the MEDLINE search (red) and number of studies included (green). **(B)** Overview of COBIDAS Domains reported in included studies. FOV, Field of view; PSF, Point spread function; ROIs, regions of interest. **(C)** Scatterplot of 141 effect sizes used in meta-analysis by published report in alphabetical order of first author. Error bars correspond to standard deviation, except for Zaaraoui et 2012 who reported range and Driver et al 2019 who reported standard error. Cb, cerebellum, DN, Dentate Nucleus; GM, grey matter; WM, white matter. Images created in R^64–66^.

We used a modified version of the checklists associated with the Committee on Best Practice in Data Analysis and Sharing (COBIDAS)^50^, appropriate to the context of ^23^Na-MRI: specifically, the sections on descriptive statistics, image acquisition reporting and pre-processing reporting. A list of reporting domains included in the modified version can be found in **Supplementary Table 4**, along with the coded results from the included reports and a summary can be seen in **Figure 3b**. Information relating to the numbers of included participants, whether informed consent was given, MRI scanner used, repetition time (TR), echo time (TE), pulse sequence and nominal resolution was provided by all included studies. No included report provided information of the distribution of handedness in the included groups. The remaining domains were reported by varying numbers of included reports.

### Descriptive

From the 28 included studies, a total of 162 effect sizes in the form of means and SD of measured TSC (mM) in healthy controls were extracted. In addition, relevant information relating to the COBIDAS domains for which all included studies had relevant data were extracted, except for ‘Scanner’ where the reporting was too variable and insufficient to permit reclassification. To account for variability in nomenclature, ‘Pulse sequence’ and ‘Tissue’ were re-coded according to **Tables 1 and Supplementary Table 5**, respectively. Tissue re-coding was informed by the wish to maximise the data per region, tissue homogeneity and physiological plausibility and minimise non-independence, and we excluded twenty effect sizes from regions too sparsely sampled to satisfy these considerations. The remaining 141 effects sizes across 14 updated Tissue regions in were taken forward for meta-analysis (**Figure 3c**): 22 effect sizes for both GM and WM regions; 14 in the Brainstem and Pons combined; 10 each in Central WM, Thalamus, and ‘GM, Temporal’ regions; eight each in ‘Deep WM’ and WM in the cerebellum and dentate nucleus (WM, Cb+DN); seven each in ‘GM, Parietal’ and Putamen ; six each in Caudate, Globus Pallidus and ‘GM, Frontal’ and five in ‘GM, Occipital’.

We also recorded the type of calibration method used - an external phantom, or internal references in the vitreous humour of the eyes or ventricles in the brain. Finally, although we only extracted data from healthy controls many of the studies in question included comparisons with individuals with a range of neurological conditions. As such, we also recorded ‘Comparison group’, in the hopes that this might capture some of the variability associated missing demographic information where controls where matched for age and sex (both incomplete COBIDAS domains, **Figure 3b**) relative to patient groups with their own typical ranges for those factors and/or the possible tendency in technical papers (without neurological comparison groups) for the data to be extracted from authors/students themselves.

#### Multi-level meta-analysis: model-fit

We used a multilevel/multivariable approach with four levels (participant, effect size, tissue regions and study), and a random effects model to pool effect sizes. The pooled mean of TSC across all 141 estimates based on the multilevel meta-analytic model was 40.51 mM (95% CI: 37.59 - 43.44; p< 0.001). We identified considerable heterogeneity (I^2^_Total=_99.4%), with the results of the standard random-effects model suggesting that most of the total variance is due to between-study heterogeneity (i.e., variance in the ‘true’ means), while the remaining (0.6%) can be attributed to sampling variance.

### Variance Components: Tissue and Study Factors

The factors Tissue and Study were included in the model to account for the nested structure of dependencies within the data: there are multiple effect sizes per paper and the effect sizes are not independent - a given individual may have contributed to multiple levels of the factor Tissue, and a given region may be estimated based on different numbers of effect sizes from multiple and varying numbers of papers. The estimated variance components were t=45.81 for the Study level, 9.93 for Tissue level and 28.96 for Effect Size level. In terms of the distribution of variance across levels as a percentage of total variance, 53.75% of the total variation in our data can be attributed to between-study heterogeneity, 11.65% to between-tissue heterogeneity and 33.99% to the effect size level (i.e. within-factor heterogeneity for Tissue), and only 0.6% is due to sampling variance. The high heterogeneity both between Studies and within Tissue, suggest that a subgroup analysis by anatomical region is appropriate.

### Model comparison: Tissue sub-group versus reduced model

In **Figure 4** we report the forest plots for anatomical regions with at least ten effect sizes only (following general statistical power guidelines on in meta-analytic sub-analyses^48^). These included (pooled mean [95% CI]): GM (45.92 mM [42.27; 49.57]), Temporal GM (45.48 mM [40.92; 50.05]), Thalamus (42.08 mM [36.38; 47.77]), Brainstem+Pons (40.99 mM [33.28;48.70]), WM (37.29 mM [33.04; 41.53]) and ‘Central’ WM (34.67 mM [27.89;41.44]). We found that the subgroup multilevel model provided a significantly better fit compared to a reduced multilevel model, as indicated by lower Akaike (AIC) and Bayesian Information Criterion (BIC). The likelihood ratio test (LRT) comparing both models is significant (χ^2^ _13_=145.50, p< 0.01). A test of the moderators ‘Tissue’ was significant, F^2^_14_ =65.34, p-val < .01), indicating that in this model the mean TSC is different between each anatomical region. However, the results indicate that high heterogeneity remains overall even with the inclusion of ‘Tissue’ level (χ^2^_127_ =9537.26, p-val < .01).

**Figure 4:**
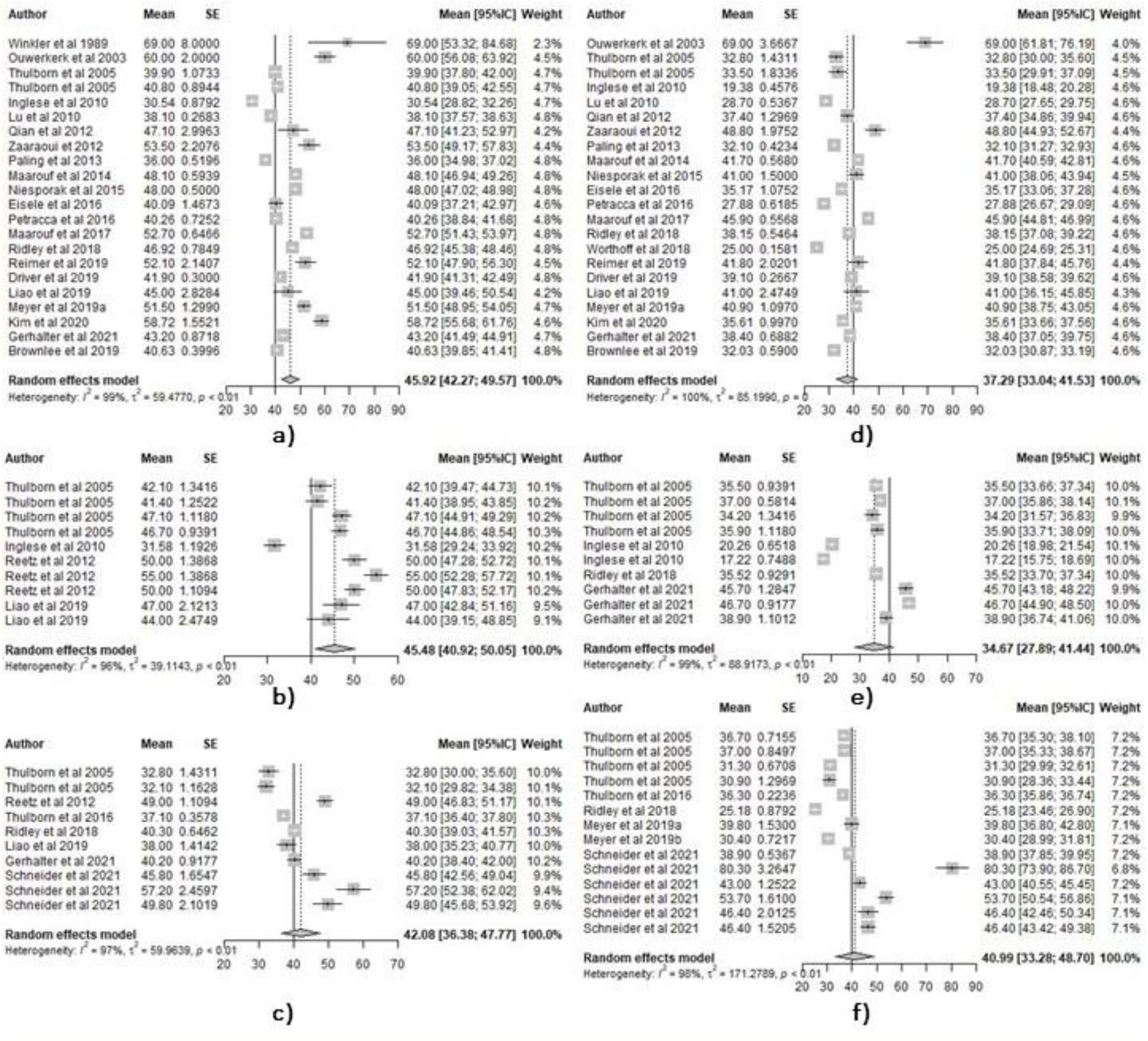
Forest plots for anatomical regions with at least ten effect sizes. a) GM, b) GM, Temporal, c) Thalamus, d) WM, e) Central WM, f) Brainstem+Pons. Each forest plot contains the effect size data, represented by grey squares scaled to their weight in the meta-analytic model and error bars corresponding to 95% confidence intervals. The regional pooled estimate for each plot is represented by a grey diamond scaled in length to the confidence interval of the pooled estimates, and a dotted reference line. The pooled overall mean of all 141 included effect sizes is represented by a solid reference line on each plot. Plots generated in R, in the metafor package^67^.

#### Precision, small study effects and publication bias

We investigated further factors impacting the distribution of results with respect to pooled means via Egger’s tests and funnel plots (**Figure 5**). The egger’s test used standard error, a measure of precision, as predictor. Things being equal, there should an inverse relationship between standard error and the probability of a given study’s estimate being different from the actual value in the population, with an expected symmetry in over- and under-estimates. Overall, the distribution of all effect sizes did not show the expected distribution, showing substantial asymmetry (Egger’s test, t=5.24, p<0.001) which remained statistically significant when outliers (identified by sensitivity analysis eliminating one by one the extreme points of the distribution) are removed (t=4.89, p<0.001). Asymmetry can be evidence for small study effects and publication bias, based on the assumption that small studies are at greatest risk of non-significant results and biasing the published literature toward the high effect sizes that are most likely to be significant with small N^48^. In our case we are investigating a measure of central tendency: mean TSC, as opposed to a standardized mean *difference*, compared to standard error. As such, we investigated the possibility that our date does not show the expected symmetry because it is constituted by sub-groups, and the ‘pooled mean’ is not the best reference given the subgroup analysis, above.

**Figure 5.**
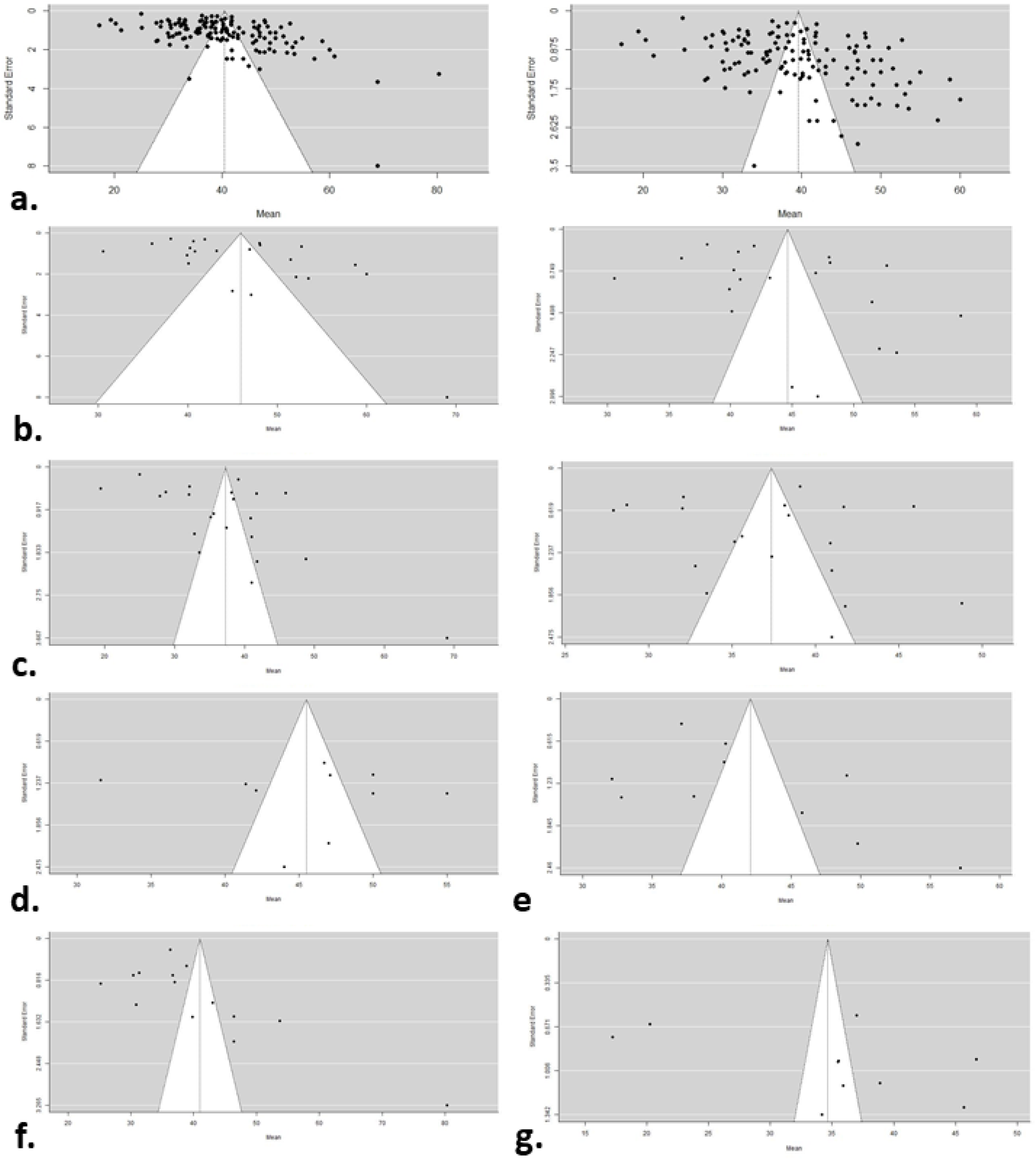
Funnel plots comparing effect sizes (mean TSC) with their precision (standard error): a) all 141 effect sizes (left) and with outliers removed (right, 136 effect sizes); b) GM effect sizes with (22, left) and without outliers (20,right); c) WM effect sizes with (22,left) and without outliers (19, right); d) GM; Temporal; e) Thalamus; f) Brainstem+Pons; g) Central WM. Images created in R^64,67^

In GM the test of asymmetry was at the threshold of significance (t=2.1, p=0.05), though after removing some outliers (60 mM^51^; 69 mM^5^) the figures become more symmetrical and the tests become non-significant (t=1.73, p=0.1) and the remaining regions do not show significant skewness. Similarly, in WM the presence of asymmetry is indicated (t=2.48, p=0.02), after removal of some outliers (19.38 mM^10^, 25 mM^57^, 69 mM^5^) distribution was no longer significantly asymmetrical. The remaining regions did not show significant asymmetry.

#### Effect of Moderators

Papers differed in their methodology across domains. We sought to understand the effect of additional methodological moderators by adding them individually and exploring their association with different mean TSC estimates. Relative to a reduced model including the factor Tissue, all additional individual factor added to the model produced a significantly better fit (**Table 2**), in terms of reducing overall heterogeneity.

**Table 2.** Tests comparing models including each methodological moderator to reduced model.

| Moderator | Likelihood ratio test |  |  | Test of residual heterogeneity |  |  |
| --- | --- | --- | --- | --- | --- | --- |
| | DF | $\chi^2$ | *p* | DF | $\chi^2$ | *p* |
| Sequence | 9 | 78.07 | < 0.01 | 118 | 4772.90 | < 0.01 |
| Comparison group | 6 | 50.73 | < 0.01 | 121 | 9081.33 | < 0.01 |
| Calibration method | 2 | 16.82 | < 0.01 | 125 | 8636.05 | < 0.01 |
| Voxel volume | 1 | 7.93 | < 0.01 | 125 | 7963.39 | < 0.01 |
| Field strength (Tesla) | 5 | 26.81 | < 0.01 | 126 | 9216.67 | < 0.01 |
| Repetition time (TR) | 1 | 29.74 | < 0.01 | 126 | 8697.18 | < 0.01 |
| Echo time (TE) | 1 | 16.01 | < 0.01 | 126 | 9529.9 | < 0.01 |

DF, degrees of freedom.

#### Moderator effects on mean TSC

Within factors, we explored the levels associated with significant differences in mean TSC between levels independent of Tissue (**Figure 6**). For studies using Sequence Type as “Radial” or “SISTINA”, the mean TSC is significantly lower than that of studies using the “DA Radial” type (*t*_*118*_=-2.95, p<0.01 and *t*_*118*_=-2, p< 0.05 respectively). Field strength of 1.5 T is associated with higher concentrations of Sodium (*t*_*122*_=2.54, p=0.01) relative to 3T. In studies with “Brain Tumour” as the Comparison Group the mean TSC is higher in controls (*t*_*118*_=2.68, p<0.01) than in studies where there was no comparison group. Tests for Calibration method, Voxel Volume, TR and TE were non-significant, indicating no association between the level of the two moderators and the mean measured sodium level, regardless of region.

**Figure 6:**
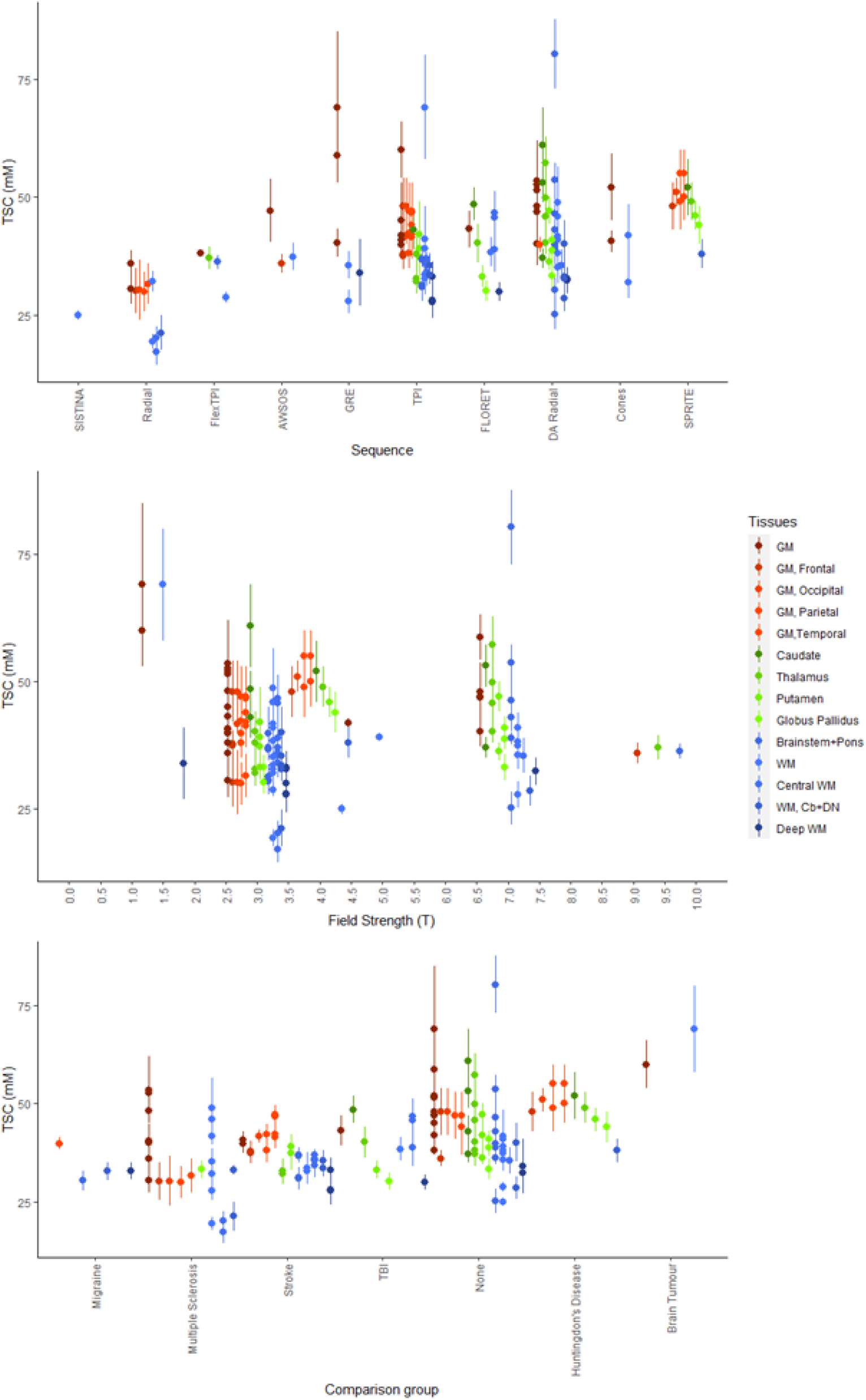
Scatterplot of 141 effect sizes used in meta-analysis ordered by moderators with a mean effect on TSC between levels independent of Tissue. Error bars correspond to standard deviation, except for Zaaraoui et 2012 (DA Radial, 3 Tesla, Multiple Sclerosis) who reported range and Driver et al 2019 (TPI, 4.7 Tesla, None) who reported standard error. Cb, cerebellum, DN, Dentate Nucleus; GM, grey matter; WM, white matter. Images created in R^64–66^.

#### Intra-regional heterogeneity

Given the high heterogeneity within the factor Tissue identified previously, we extended our analysis to identifying where inclusion of methodological moderators reduces the heterogeneity of estimates within anatomical regions (with at least 10 effects sizes) suggesting the pooled estimates in the reduced model are impacted by differences in a given factor. We compared the specific heterogeneity (tau) of a region in the reduced model compared to the model with the methodological factor using Hedges’ g, identifying ‘significant’ reductions in the form of a standardized mean difference whose 95% CIs did not cross zero (**Supplementary Table 6, Supplementary Figure 1**).

‘Sequence’ as moderator reduces the heterogeneity for GM, GM-Temporal and WM regions. Assuming “Comparison Group” reduces heterogeneity in GM, Brainstem+Pons, and WM. “Calibration method” reduces heterogeneity within GM. Residual heterogeneity is reduced in “Brainstem+Pons” when “Voxel Volume” is used as a moderator. “TR” reduced residual heterogeneity when used as a moderator in Temporal GM and the thalamus. “TE” reduced heterogeneity in Temporal GM and Central WM. Field strength did not contribute to explaining the variability within anatomical regions. Note that for all methodological factors, the test of residual heterogeneity for the model overall remained significant (**Table 2**)

## Discussion

Data from 28 studies, identified by literature search, were explored via meta-analysis - to our knowledge the first such attempt in the context of data from ^23^Na-MRI. The overall pooled estimate from all 141 across all 28 studies was 40.51 mM (95% CI: 37.59 - 43.44), well within the ranges suggested by parenchymal volume models (37-45mM^6, 12,20^). Meta-analytic estimates were associated with high heterogeneity, which further analysis suggested was largely associated with between-study heterogeneity. This supports the idea that there is underlying differences in the ‘true means’ the different studies are trying to measure – and that a parenchymal estimate is not sufficient to characterise the range of empirical values obtained from different brain regions. Pooled estimates based on the extant literature of TSC estimates in human samples are 45.92 mM [42.27; 49.57] for GM and 37.29 mM [33.04; 41.53] for WM. This is noticeably higher than tissue volume model-based estimates for both tissue types, with examples including estimates of 20-33mM in WM and for 30-35 for GM^38,68^.

A number of potential sources may account for discrepancy between theoretical tissue volume models and the empirical estimates. One is partial volume effects, and other methodological factors impacting acquisition of the estimates making up the literature explored here. The models use simplifying assumptions, most notably that they have adequately captured the relevant influences on sodium ^23^Na-MRI measurement with a limited number of compartmental volume contributions, and that these compartments are internally homogenous. For example, the extracellular compartment is generally (but not always^40^) considered to include everything outside cells membranes, thereby subsuming the interstitial extracellular matrix^69^ and vascular spaces and attributing the ^23^Na concentration of ‘pure’ CSF to the entire compartment ^20,38,40,70^. Similarly, the contribution of membranes, lipids like myelin and other ‘solids’ are assumed to be captured by a single volume contribution with no sodium contribution (on the basis they exclude sodium and should reduce overall measured signal for a given volume^40,70^), and can be summarised by a single fractional variable (e.g. 0.7-0.9^38,40,42^) for a given tissue type. It would be interesting to see how the modification of any or all these impacts interact to produce expected values and how this might be applied to regional estimates.

More granular tissue models are currently lacking, as the requisite cellular data is not available. A potentially relevant factor is regional variation in the ratios of different cell types in the context of divergent sodium concentrations, for example astrocytes have approximately twice the cytosolic sodium concentration (∼15-20mM) compared to neurons (∼10mM) in rodent samples^71–75^. Recent automated immunocytochemical techniques have provided much needed information, including correcting widespread misconceptions about neuronal versus non-neuronal populations and masses, precise data on cell volumes and their variation – which would be relevant for building regionally specific volume models for ^23^Na-MRI – are not yet available^43,44^. In the absence of complete histologic information, another source of insight could come from comparing ^23^Na-MRI to other imaging indices that might capture relevant features, with other quantitative imaging modalities being of particular interest. Parallel changes in ^23^Na-MRI measures and diffusion imaging^15,76–78^ and proton density^79^ are suggestive, but extending these to direct evaluations of the redundancy and complementarity with other quantitative modalities^63^, especially in healthy controls, in a range of regions and tissue structures would be a welcome development.

While the empirical values are higher than the model-based estimates for a given tissue, the relative values of different tissue’s concentrations (e.g. GM>WM) is preserved. The difference in myelin content between grey and white matter – captured in the differences in ‘solid fractions that are usually assigned – may account for some of this difference. Indeed, among the subcortical regions we were able to provide pooled estimates for (>10 effect sizes), it is noticeable that intermediate values were described. The ostensibly GM but highly myelinated Thalamus has a lower value (42.08 mM [36.38; 47.77]) than some other GM estimates (45.48 mM [40.92; 50.05] for Temporal, GM), while ROIs sampling regions that are likely to be predominantly WM but with contributions from GM nuclei like the brainstem and pons indicate higher values (40.99 mM [33.28;48.70]) than some other regions (34.67 mM [27.89;41.44] for ‘Central WM’). While the overlap between confidence intervals and the remaining heterogeneity limits the certainty of these precise pooled means, which should not be taken as definitive given the limitations of the available literature as represented in this meta-analysis, the importance of considering the variation in apparent ^23^Na associated with regional differences is reflected in the significant improvement in the model when including the tissue factor and the finding of significant differences in mean TSC between levels/regions.

Given the remaining unexplained heterogeneity even after the Tissue factor was involved, we explored additional methodological factors where full reporting made this possible. The fundamentally most limiting property of ^23^Na-MRI is the reduced nuclear MR sensitivity and relative abundance of sodium and other non-^1^H based contrasts^80^, leading to reduced signal to noise ratios and resolution, and consequently to partial volume effects. A given ‘Sequence’ is an attempt mitigate between trade-offs in acquisition parameters with impacts on available signal and resolution (eg. TE, TR, Voxel Volume, Field Strength). Different internal and external calibration methods provide scope for different degrees of experimenter error as well as spatial and physiological variability^81^. Comparison group may reflect demographic factors of potential relevance^82^ reflecting the target patient group and other variable experimenter/location factors. We were unable to consider the majority of methodological and demographic factors sought by the modified COBIDAS checklist because information was not reported (**Figure 3b**). Explicit reference to and information pertaining to the following COBIDAS domain were not identified in for age, handedness, sex, coil information, acquisition time, processing software used, flip angle, segmentation and ROI definition, details of normalisation / registration, field of view, information pertaining to smoothing or point spread function.

We found each fully-reported methodological factor with included beyond tissue improved the meta-analytic model fit (**Table 2**), but that significant residual heterogeneity remained regardless. Only Field Strength, Sequence, and Comparison Group differed in mean TSC between specific levels independently of tissue. It is possible these specific differences may be driven by sampling issues: a single study is sampled for the SISTINA level of the factor Sequence while another single study was sampled both at the 1.5 Tesla level of the factor Field Strength and the Brain Tumour level of Comparison Group, which was also identified as an outlier in the assessment of asymmetry in standard error distribution for both GM and WM^5^ (**Figure 5, Figure 6**). Considered in combination with Tissue, all methodological factors except for Field Strength reduced heterogeneity in some regions when included (**Supplementary Figure 1)**. Collectively these results stress the importance of methodological factors but also the limitations of the available literature and underline the need for more and completely reported data covering multiple acquisition schemes and brain regions.

We analysed estimates of Total/Tissue Sodium Concentration, as the most common measure available. Other ^23^Na-MRI derived metrics are possible, for example there are approaches that measure^56^ or filter sodium signal based on relaxation behaviour (e.g. inversion recovery, IR^83^), or multiple quantum filtering (MQF)^16^. I*n principle*, any *specific* measurement of in vivo sodium by ^23^Na-MRI – TSC, IR, MQF or other – cannot be said to derive from a single cellular-level tissue compartment^84^. However, while attribution to different sources is a subject of longstanding and ongoing investigation^3,32,85^, it is legitimate to discuss a *difference or change* in measured parameters (between conditions and across spatial/temporal domains) in terms of changes in concentration or structure in sub-compartments that may have contributed, even when the latter are below the limit of resolution. In practice, precisely attributing changes in empirical MR-level estimates to compartmental micro-features is unlikely to be definitive because these factors rarely alter in isolation. For example, while pathological TSC alterations may be related to metabolic impairments of transmembrane ^23^Na exchange, they may also reflect changes in cellular death, swelling, proliferation etc^86–88^. Fundamentally, ^23^Na-MRI appears to be a more sensitive than specific measure, and a claim that it is sensitive to variation in a particular structural, functional or patho-/physiological context is an empirical question to answered by further appropriate data and not modelling or a priori arguments from incomplete biophysical data alone.

Some further limitations should be noted and considered in future analysis. Variable reporting led required accommodations be made in several factors that were included in the analysis. Allocation of a given data point to a particular level of Tissue was based on explicit textual references in the included studies, but the method of segmentation, precise anatomic boundaries, use of atlases and coordinates where not always clear. This may be another source of heterogeneity in the results, and again highlights the need for clear reporting. We also considered only published reports, and not ‘grey literature’ (e.g. dissertations, preprints, government reports, or conference proceedings)^48^ which might could potentially improve sampling. We produced and examined a central tendency measure via mean estimates of TSC, but if and when sufficient data are available meta-analytic analysis could be applied to other ^23^Na-MRI metrics as well as combined estimates of differences between groups, structures and states.

## Conclusions

Data from 28 studies, identified by literature search, were explored via meta-analysis – to our knowledge the first such attempt in the context of data from ^23^Na-MRI. The nested nature of the data, due in part to accommodations made to the variability of reporting in the published studies, lead to the use of a multi-level meta-analytic approach. We produced pooled meta-analytic estimates of brain TSC, but significant remaining heterogeneity limits the certainty and precision associated with the estimates. Consideration of tissue explains part of that heterogeneity, but not all. Where full reporting permitted, methodological moderators were explored. While inclusion reduces heterogeneity within certain tissue regions, and effects measured TSC levels, substantial residual heterogeneity remains. The current estimates provide an empirical point of departure for better understanding of variability in ^23^Na-MRI. Improving on current estimates supports: (1) larger, more representative data collection/sharing, including (2) regional data, and (3) agreement on full reporting standards.

## Methods

### Literature search

The following MEDLINE search was run by BR via pubmed.ncbi.nlm.nih.gov on 12/07/2021: “Brain” [Title/Abstract] AND ((((sodium MRI [Title/Abstract]) OR 23Na MRI [Title/Abstract]) OR sodium imaging [Title/Abstract]) OR 23Na imaging [Title/Abstract])”. Bibliographies of potentially eligible studies were consulted and studies of potential relevance to ^23^Na-MRI were included in the screening.

Recovered records were excluded based on the abstract or full text if they were non-experimental, non-original report (review/commentary), conference proceedings, phantom-only studies, concerned with cultured tissue or organs other than the brain, non-human subjects concerning organs other than the brain, did not include estimates of sodium concentrations determined by quantitative ^23^Na-MRI in healthy subjects without known neurological conditions. Studies considering only estimates in overall parenchyma, or where it was not possible to attribute estates to a specified anatomical region were also excluded. Screening and full text review were performed by BR with reference to other authors as necessary.

### Data Extraction

Initial extraction of data pertaining to ^23^Na-MRI concentrations and methodological domains was performed by BR, with verification and consultation with WZ. Data re-coding, as discussed in the “descriptive” section in Results, was based on consensus decisions by WZ/BR for “Sequence” and FN/BR for “Tissue” (See **Tables 1 and Supplementary Table 5)**. Where a given study included multiple potentially relevant samples a context-based decision was made: where the same individuals were sampled differing methods we took the highest field / lowest resolution observation for Qian et al^53^; where the same individual was sampled multiple times with the same methods (reproducibility studies) we took the overall mean across samples for Riemer et al^58^ and Meyer et al^24^; where PVC-corrected values by different methods were used in Kim et al^37^ we used the spill-over and ventricular CSF-based PVC-corrected values. Where multiple TEs where reported we took the minimum value.

### Meta-analysis

Meta-analyses were conducted in R (**R-4**.**1**.**2)** using the “metafor” package^67^. The restricted maximum likelihood estimator^89^ was used to calculate the heterogeneity variance (τ2) and we used Knapp-Hartung adjustments^90^ (Knapp & Hartung, 2003) to calculate the confidence interval around the pooled effect. Multi-level models were investigated to account for any correlations induced by the multi-level structure of the data. To account for correlated sampling errors due to different effect sizes being based on the same sample of patients; to take this into account, we used a Correlated and Hierarchical Effects (CHE) model, an extension of the multilevel model that considers the correlation of effect sizes within clusters, in this case the factor ‘Paper’. A robust Sandwich covariate estimator was used to confidence intervals and relative p-values. Egger’s tests were used to evaluate asymmetry of funnel plots, based on weighted regression models with multiplicative dispersion, with standard error as the predictor.

## Author contributions

BR conceived and planned the work, designed, and ran the literature search. BR, WZ, and FN contributed to data extraction. BR and FM designed and ran the analysis and prepared figures. All authors contributed to interpretation of the results. BR drafted the manuscript, which was revised by all authors, who agreed to the final version.

## Data availability statement

All data generated or analysed during this study are included in this published article and its Supplementary Information files.

## Additional Information

The authors declare no competing interests.

