## Supplementary Figure 1 for "Variability by region and method in human brain sodium concentrations estimated by ^23^Na magnetic resonance imaging: a meta-analysis"

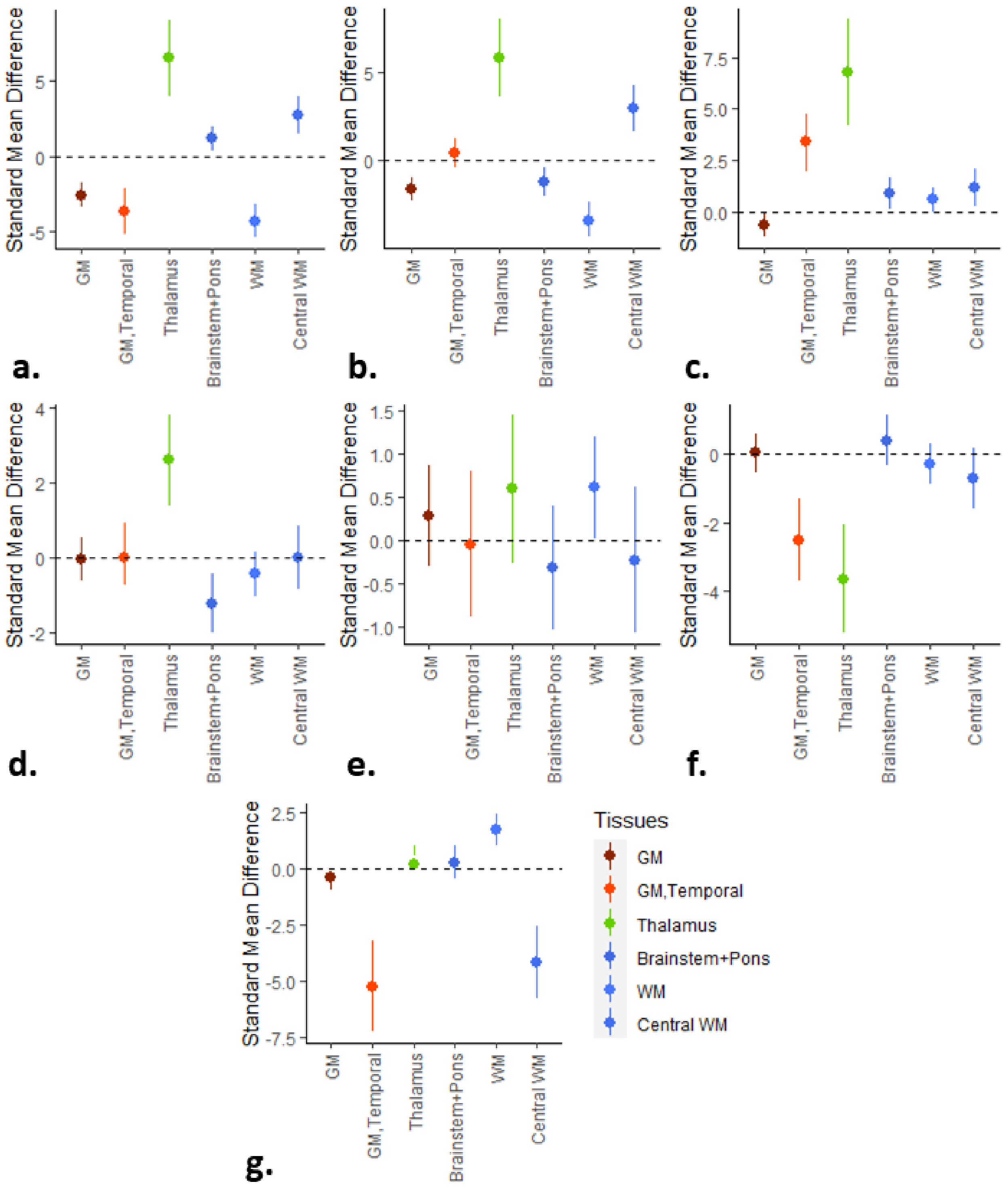


**Supplementary Figure 1: Scatterplots of standardized mean differences (Hedges' g) in regional heterogeneity (tau) between models with and without methodological moderators.** a) Sequence, b) Comparison group, c) Calibration method, d) Voxel volume, e) Field Strength, f) TR, g) TE. Error bars correspond to 95% confidence intervals. Images created in R[64–66]
