## Supplementary tables for "Variability by region and method in human brain sodium concentrations estimated by ^23^Na magnetic resonance imaging: a meta-analysis"

### Supplementary Table 1: PRISMA 2020 checklist

| Section and Topic | Item # | Checklist item | Location where item is reported |
|---|---|---|---|
| **TITLE** | | | |
| Title | 1 | Identify the report as a systematic review. | N/A |
| **ABSTRACT** | | | |
| Abstract | 2 | See the PRISMA 2020 for Abstracts checklist. | Supplementary Table 2 |
| **INTRODUCTION** | | | |
| Rationale | 3 | Describe the rationale for the review in the context of existing knowledge. | Introduction |
| Objectives | 4 | Provide an explicit statement of the objective(s) or question(s) the review addresses. | Introduction |
| **METHODS** | | | |
| Eligibility criteria | 5 | Specify the inclusion and exclusion criteria for the review and how studies were grouped for the syntheses. | Methods/Results |
| Information sources | 6 | Specify all databases, registers, websites, organisations, reference lists and other sources searched or consulted to identify studies. Specify the date when each source was last searched or consulted. | Methods/Results |
| Search strategy | 7 | Present the full search strategies for all databases, registers and websites, including any filters and limits used. | Methods |
| Selection process | 8 | Specify the methods used to decide whether a study met the inclusion criteria of the review, including how many reviewers screened each record and each report retrieved, whether they worked independently, and if applicable, details of automation tools used in the process. | Methods |
| Data collection process | 9 | Specify the methods used to collect data from reports, including how many reviewers collected data from each report, whether they worked independently, any processes for obtaining or confirming data from study investigators, and if applicable, details of automation tools used in the process. | Methods |
| Data items | 10a | List and define all outcomes for which data were sought. Specify whether all results that were compatible with each outcome domain in each study were sought (e.g. for all measures, time points, analyses), and if not, the methods used to decide which results to collect. | Methods |
| | 10b | List and define all other variables for which data were sought (e.g. participant and intervention characteristics, funding sources). Describe any assumptions made about any missing or unclear information. | Methods |
| Study risk of bias assessment | 11 | Specify the methods used to assess risk of bias in the included studies, including details of the tool(s) used, how many reviewers assessed each study and whether they worked independently, and if applicable, details of automation tools used in the process. | N/A – see COBIDAS checklist & results/discussion |
| Effect measures | 12 | Specify for each outcome the effect measure(s) (e.g. risk ratio, mean difference) used in the synthesis or presentation of results. | Methods/Results |
| Synthesis methods | 13a | Describe the processes used to decide which studies were eligible for each synthesis (e.g. tabulating the study intervention characteristics and comparing against the planned groups for each synthesis (item #5)). | Methods/Results |
| | 13b | Describe any methods required to prepare the data for presentation or synthesis, such as handling of missing summary statistics, or data conversions. | Methods/Results |

| Section and Topic | Item # | Checklist item | Location where item is reported |
|---|---|---|---|
| | 13c | Describe any methods used to tabulate or visually display results of individual studies and syntheses. | Methods/Results |
| | 13d | Describe any methods used to synthesize results and provide a rationale for the choice(s). If meta-analysis was performed, describe the model(s), method(s) to identify the presence and extent of statistical heterogeneity, and software package(s) used. | Methods/Results |
| | 13e | Describe any methods used to explore possible causes of heterogeneity among study results (e.g. subgroup analysis, meta-regression). | Methods/Results |
| | 13f | Describe any sensitivity analyses conducted to assess robustness of the synthesized results. | Methods/Results |
| Reporting bias assessment | 14 | Describe any methods used to assess risk of bias due to missing results in a synthesis (arising from reporting biases). | N/A – see COBIDAS checklist & results/discussion |
| Certainty assessment | 15 | Describe any methods used to assess certainty (or confidence) in the body of evidence for an outcome. | N/A – see COBIDAS checklist & results/discussion |
| **RESULTS** | | | |
| Study selection | 16a | Describe the results of the search and selection process, from the number of records identified in the search to the number of studies included in the review, ideally using a flow diagram. | Results, Search Results |
| | 16b | Cite studies that might appear to meet the inclusion criteria, but which were excluded, and explain why they were excluded. | Results |
| Study characteristics | 17 | Cite each included study and present its characteristics. | Results, Included study |
| Risk of bias in studies | 18 | Present assessments of risk of bias for each included study. | N/A – see COBIDAS checklist & results/discussion |
| Results of individual studies | 19 | For all outcomes, present, for each study: (a) summary statistics for each group (where appropriate) and (b) an effect estimate and its precision (e.g. confidence/credible interval), ideally using structured tables or plots. | Results / Materials and Data |
| Results of syntheses | 20a | For each synthesis, briefly summarise the characteristics and risk of bias among contributing studies. | N/A – see COBIDAS checklist & results/discussion |
| | 20b | Present results of all statistical syntheses conducted. If meta-analysis was done, present for each the summary estimate and its precision (e.g. confidence/credible interval) and measures of statistical heterogeneity. If comparing groups, describe the direction of the effect. | Results, Meta-analysis |
| | 20c | Present results of all investigations of possible causes of heterogeneity among study results. | Results, Meta-analysis |
| | 20d | Present results of all sensitivity analyses conducted to assess the robustness of the synthesized results. | Results, Meta-analysis |
| Reporting biases | 21 | Present assessments of risk of bias due to missing results (arising from reporting biases) for each synthesis assessed. | N/A – see COBIDAS checklist & results/discussion |
| Certainty of evidence | 22 | Present assessments of certainty (or confidence) in the body of evidence for each outcome assessed. | N/A – see COBIDAS checklist & results/discussion |
| **DISCUSSION** | | | |
| Discussion | 23a | Provide a general interpretation of the results in the context of other evidence. | Discussion |
| | 23b | Discuss any limitations of the evidence included in the review. | Discussion |
| | 23c | Discuss any limitations of the review processes used. | Discussion |

| Section and Topic | Item # | Checklist item | Location where item is reported |
|---|---|---|---|
| | 23d | Discuss implications of the results for practice, policy, and future research. | Discussion |
| **OTHER INFORMATION** | | | |
| Registration and protocol | 24a | Provide registration information for the review, including register name and registration number, or state that the review was not registered. | N/A |
| | 24b | Indicate where the review protocol can be accessed, or state that a protocol was not prepared. | N/A |
| | 24c | Describe and explain any amendments to information provided at registration or in the protocol. | N/A |
| Support | 25 | Describe sources of financial or non-financial support for the review, and the role of the funders or sponsors in the review. | Sources of support |
| Competing interests | 26 | Declare any competing interests of review authors. | Conflicts of Interest |
| Availability of data, code and other materials | 27 | Report which of the following are publicly available and where they can be found: template data collection forms; data extracted from included studies; data used for all analyses; analytic code; any other materials used in the review. | Materials and Data |

*From:* Page MJ, McKenzie JE, Bossuyt PM, Boutron I, Hoffmann TC, Mulrow CD, et al. The PRISMA 2020 statement: an updated guideline for reporting systematic reviews. BMJ 2021;372:n71. doi: 10.1136/bmj.n71

For more information, visit: http://www.prisma-statement.org/

#### Supplementary Table 2: PRISMA 2020 abstract checklist

| Section and Topic | Item # | Checklist item | Reported (Yes/No) |
|---|---|---|---|
| **TITLE** | | | |
| Title | 1 | Identify the report as a systematic review. | NA |
| **BACKGROUND** | | | |
| Objectives | 2 | Provide an explicit statement of the main objective(s) or question(s) the review addresses. | Yes |
| **METHODS** | | | |
| Eligibility criteria | 3 | Specify the inclusion and exclusion criteria for the review. | Yes |
| Information sources | 4 | Specify the information sources (e.g. databases, registers) used to identify studies and the date when each was last searched. | Yes |
| Risk of bias | 5 | Specify the methods used to assess risk of bias in the included studies. | NA |
| Synthesis of results | 6 | Specify the methods used to present and synthesise results. | Yes |
| **RESULTS** | | | |
| Included studies | 7 | Give the total number of included studies and participants and summarise relevant characteristics of studies. | Yes |
| Synthesis of results | 8 | Present results for main outcomes, preferably indicating the number of included studies and participants for each. If meta-analysis was done, report the summary estimate and confidence/credible interval. If comparing groups, indicate the direction of the effect (i.e. which group is favoured). | Yes |
| **DISCUSSION** | | | |
| Limitations of evidence | 9 | Provide a brief summary of the limitations of the evidence included in the review (e.g. study risk of bias, inconsistency and imprecision). | Yes |
| Interpretation | 10 | Provide a general interpretation of the results and important implications. | Yes |
| **OTHER** | | | |
| Funding | 11 | Specify the primary source of funding for the review. | NA |
| Registration | 12 | Provide the register name and registration number. | NA |

*From:* Page MJ, McKenzie JE, Bossuyt PM, Boutron I, Hoffmann TC, Mulrow CD, et al. The PRISMA 2020 statement: an updated guideline for reporting systematic reviews. BMJ 2021;372:n71. doi: 10.1136/bmj.n71

#### Supplementary Table 3: All excluded studies with reasons for rejection

| **Paper** | **Details** | **Notes** | **Reason for rejection** |
|---|---|---|---|
| Abad et al 2018 | Pain. 2018 October ; 159(10): 2058–2065 | Rat, Migraine | Animal model |
| Adlung et al 2021 | NMR in Biomedicine. 2021;e4474. | Human, Brain, Stroke | no controls |
| Adlung et al 2021 | Cerebrovasc Dis. 2021;50(3):347-355. | Human, Brain, Stroke | no controls |
| Albert et al 1993 | NMR Biomed. 1993 Jan-Feb;6(1):7-20. | Blood, Rat, shift reagent | organ other than brain |
| Alecci et al 2006 | J Magn Reson. 2006 Aug;181(2):203-11. | Coil design | no concentration |
| Allis et al 1991 | JMR 93, 7 l-76 ( 199 1) | Cardiac, Rat, TQF | organ other than brain |
| Atkinson & Thulborn 2010 | NeuroImage 51 (2010) 723–733 | Brain, Human | no concentration |
| Atkinson et al 2007 | J Magn Reson Imaging. 2007 Nov;26(5):1222-7. | Human, UHF safety study | no concentration |
| Atkinson et al 2011 | MRM 66:1089–1099 (2011) | Human, Brain | Parenchyma/ anatomical attribution not possible |
| Atkinson et al 2012 | Magnetic Resonance in Medicine 68:751–761 (2012) | Methodology, coregistration / longitudinal data | no concentration (in usable form) |
| Atkinson et al 2013 | Magn Reson Med. 2014 May;71(5):1819-25 | Brain, Human, UHF, 39P /23Na | no concentration |
| Atthe et al 2009 | Am J Physiol Renal Physiol. 2009 Nov;297(5):F1288-98 | Kidney, Rat | organ other than brain |
| Aursand et al 2009 | J Agric Food Chem. 2009 Jan 14;57(1):46-54 | Salmon | organ other than brain |
| Babsky et al 2006 | J Magn Reson Imaging. 2006 Jul;24(1):132-9. | Rat, Glioma | Animal model, no concentration |
| Babsky et al 2007 | Magn Reson Imaging 2007;25: 1015–1023 | Tumour, Rat, TQF | organ other than brain |
| Babsky et al 2008 | Magn Reson Med. 2008 Mar;59(3):485-91. | Muscle, Rat | organ other than brain |
| Baier et al 2014 | Magn Reson Mater Phy (2014) 27:71–79 | Rat, Brain, Stroke | Animal model, no concentration |
| Balschi et al 1990 | NMR Biomed. 1990 Apr;3(2):47-58. | Muscle, Rat | organ other than brain |
| Bansal & Seshan 1995 | J Magn Reson Imaging. 1995 Nov-Dec;5(6):761-7 | Kidney, Rabbit | organ other than brain |
| Bansal 1994 | MR pulses 1994: 1;18 | No access | No access |
| Bansal et al 1992 | JMRI 1992 2:385-391 | Rat, Brain | Animal model, no concentration |
| Bansal et al 1993 | Biochemisty 1993; 32: 5638-5643 | Liver, Rat | organ other than brain |
| Bansal et al 2000 | Magn Reson Med 11: 532-538 | Muscle, Human | organ other than brain |
| Bárány & Venkatasubramanian 1986 | Physiol Chem Phys Med NMR. 1986;18(4):233-41. | Capillary, rat | organ other than brain |
| Barfuss et al 1988 | Radiology. 1988 Dec;169(3):811-6. | signal intensity | no concentration |
| Bartha & Menon | Magn Reson Med 52:407–410 (2004) | Human, T2 relaxation, long component only | no concentration |

| | | | |
|---|---|---|---|
| Bartha et al 2004 | MRI 22 (2004) 983–991 | Rat, Brain, Stroke | Animal model, no concentration |
| Bartha et al 2008 | AJNR Am J Neuroradiol. 2008 Mar;29(3):464-70. | Human, Glioma | no concentration |
| Behl et al 2015 | Magnetic Resonance in Medicine 2015 75(4) | Brain, Human, Compressed sensing | no concentration |
| Benkhedah et al 2013 | Magnetic Resonance in Medicine 70:754–765 (2013) | Sequence development, Biexponential-weighted imaging | no concentration |
| Benkhedah et al 2014 | Journal of Magnetic Resonance 240 (2014) 67–76 | Sequence development, Biexponential-weighted imaging | no concentration |
| Benkhedah et al 2016 | Magnetic Resonance in Medicine 75:527–536 (2016) | Coil development | no concentration |
| Berendsen & Edzes 1973 | Ann N Y Acad Sci. 1973 Mar 30;204:459-85. | Muscle, Rat | organ other than brain |
| Bertram et al 2005 | J Agric Food Chem. 2005 Oct 5;53(20):7814-8. | Cured meat | organ other than brain |
| Betz et al 1994 | J Cereb Blood Flow Metab. 1994 Jan;14(1):29-37 | Rat, Brain, Stroke | Animal model, no concentration |
| Biller et al 2016 | Sci Rep. 2016 Aug 10;6:31269. | Brain, Human, Multiple sclerosis | no control concentration |
| Biller et al 2016 | AJNR 2016, 37 (1) 66-73 | Human, Brain, Tumour | no control concentration |
| Blomer et al 2021 | Curr Protoc. 2021 Mar;1(3):e64 | Mouse, Ultrafast Sodium imaging | cultured tissue / in vitro |
| Blum et al 1988 | Am J Physiol. 1988 Sep;255(3 Pt 1):C377-84. | muscle, Rat | organ other than brain |
| Blum et al 1991 | Magn Reson Med. 1991 Apr;18(2):348-57. | Liver, Rat | organ other than brain |
| Blunck et al 2018 | Magn Reson Med. 2018 Apr;79(4):1950-1961 | Brain, Human, T2 | no concentration |
| Blunck et al 2018 | Magn Reson Med. 2019 Feb;81(2):1172-1180 | Sequence development (zGRF-RHE) | no concentration |
| Blunck et al 2019 | Magn Reson Med. 2019;00:1–9. | Compressed sensing, Concentrations not in usable form | no concentration (in usable form) |
| Boada et al 1994 | MRM 32:219-223 (1994) | Rat | Parenchyma/ anatomical attribution not possible |
| Boada et al 1997 | Int. J. Imaging Systems Technol. 8, 544–550 | Human, Brain | Parenchyma/ anatomical attribution not possible |
| Boada et al 1997 | Magn Reson Med. 1997 May;37(5):706-15. | Sequence development (TPI) | no concentration |
| Boada et al 1997 | Magn Reson Med. 1997 Dec;38(6):1022-8. | Sequence development (TPI) | no concentration |
| Boada et al 1997 | Magn Reson Med. 1997 Mar;37(3):470-7. | Sequence development (TPI) | no concentration |
| Boada et al 2004 | Conf Proc IEEE Eng Med Biol Soc. 2004;7:5238-41. | Sequence development / application (TQF) | no concentration |
| Boada et al 2005 | Conf Proc IEEE Eng Med Biol Soc. 2005;1:731- | Macaque, Brain, Stroke | Animal model, no concentration |

| | | | |
|---|---|---|---|
| Boada et al 2005 | Curr Top Dev Biol. 2005;70:77-101 | Macaque, Brain, Stroke | Animal model, no concentration (in non-figure form) |
| Boada et al 2012 | Transl Stroke Res. 2012 Jun;3(2):236-45 | Review | Review/non-experimental |
| Boada et al 2012 | Transl Stroke Res. 2012 Jun;3(2):236-45 | Review, Stroke | Review/non-experimental |
| Borthakur et al 1999 | J Magn Reson 1999;141:286–90 | Cartilage, human | organ other than brain |
| Borthakur et al 2002 | Radiology. 2002 Aug;224(2):598-602. | Wrist Joint | organ other than brain |
| Bothakur et al 2006 | NMR Biomed 2006;19:781–821 | Cartilage, review | Review/non-experimental |
| Bottomley 2007 | eMagRes. 2007 | Review | Review/non-experimental |
| Boulanger & Vinay 1989 | Can J Physiol Pharmacol. 1989 Aug;67(8):820-8 | Review | Review/non-experimental |
| Bull 1969 | J Magn Reson (1969), 8, (4): 344-353 | Review/theory , Quadrupolar relaxation | Review/non-experimental |
| Burnstein & Springer 2019 | Magn Reson Med. 2019;82:521–524. | Review / Letter, 23Na interpretation | Review/non-experimental |
| Burstein & Fossel 1987 | Magn Reson Med. 1987 Mar;4(3):261-73. | Cardiac, Frog | organ other than brain |
| Burstein & Mattingly 1989 | J Magn Reson 1989:83:187–204. | Cardiac | organ other than brain |
| Burstein et al 2009 | Radiol Clin North Am. 2009 Jul;47(4):675-86 | Cartilage | organ other than brain |
| Buster et al 1990 | Magn Reson Med. 1990 Jul;15(1):25-32. | Cardiac, Rat, Shift Reagent | organ other than brain |
| Butwell et al 1991 | Invest Radiol. 1991 Dec;26(12):1079-82. | Cardiac, Rat | organ other than brain |
| Bydder et al 2019 | NeuroImage 184 (2019) 771–780 | Brain, Human, Dynamic Na-MRI, Task-Based | Parenchyma/ anatomical attribution not possible |
| Cannon et al 1986 | J Am Coll Cardiol. 1986 Mar;7(3):573-9. | Cardiac, dog | organ other than brain |
| Chang et al 2010 | Eur Radiol. 2010 Aug;20(8):2039-46 | Muscle, Human | organ other than brain |
| Chang et al 2010 | J Anat. 2010 Dec;217(6):694-704 | Rat, Na channel expression | cultured tissue / in vitro |
| Chang et al 2012 | Eur Radiol. 2012 Jun;22(6):1341-9 | Cartilage, Human | organ other than brain |
| Choi & Gold 2011 | Magn Reson Imaging Clin N Am. 2011 May;19(2):249-82 | Cartilage | organ other than brain |
| Christensen et al 1996 | MRM 36:83-9 (1996) | Rat | Parenchyma/ anatomical attribution not possible |
| Chu et al 1990 | Magn Reson Med. 1990 Feb;13(2):239-62. | Phantom, Shift Reagent | Phantom |
| Chung et al 1990 | J. Magn. Res., 1990. 88: p. 440-447 | No access | No access |
| Clayton et al 2003 | Acad Radiol 2003; 10:358–365 | Sequence development | no concentration |
| Colet et al 1998 | Magn Reson Med 1998; 39: 155-159 | Liver, Mouse | organ other than brain |
| Collorone et al 2021 | *Brain*, awab043 | Human, Brain, Multiple Sclerosis | no concentration |

| | | | |
|---|---|---|---|
| Constantinides et al 2000 | Magn Reson Imaging. 2000 May;18(4):461-71. | Cardiac, canine | organ other than brain |
| Constantinides et al 2000 | Radiology. 2000 Aug;216(2):559-68. | Muscle | organ other than brain |
| Constantinides et al 2001 | Magn Reson Med. 2001 Dec;46(6):1144-51. | Cardiac, Dog | organ other than brain |
| Constantinides et al 2001 | Magn Reson Med. 2001 Dec;46(6):1164-8. | Cardiac, Dog | organ other than brain |
| Coste et al 2019 | MRI Volume 58, May 2019, Pages 116-124 | Human, Brain | Parenchyma/ anatomical attribution not possible |
| Crema et al 2011 | Radiographics. 2011 Jan-Feb;31(1):37-61. | Cartilage | organ other than brain |
| Donandieu et al 2017 | Mult Scler. 2019 Jan;25(1):39-47 | Human, Brain, Multiple Sclerosis | no concentration |
| Eisele et al 2019 | Multiple Sclerosis and Related Disorders 29 (2019) 48–54 | Brain, Human, Multiple sclerosis | no control concentration |
| Eleff et al 1993 | Magn Reson Med. 1993 Jul;30(1):11-7 | Brain, dog, shift reagents | no concentration |
| Eliav et al 1992 | Journal of Magnetic Resonance (1969) 98(1):223–229 | Bovine cartilage, DQF, TQF | organ other than brain |
| Eliav et al 2013 | JMR 231 (2013) 61–65 | Bovine optic nerve | organ other than brain |
| Endre et al 1989 | Magn Reson Med. 1989 Aug;11(2):267-74. | Kidney, Rat | organ other than brain |
| Feinberg et al 1985 | Radiology. 1985 Jul;156(1):133-8. | signal intensity | no concentration |
| Felix et al 2020 | Front Physiol . 2020 Aug 12;11:871 | Review, Astroglia | Review/non-experimental |
| Fiege et al 2013 | Magn Reson Med. 2013 Jun;69(6):1691-6 | Sequence development, TSF, SISTINA | no concentration |
| Filipis et al 2020 | J Physiol. 2021 Jan;599(1):49-66 | mouse, brain section | cultured tissue / in vitro |
| Fleysher et al 2009 | Magn Reson Med. 62:1338–1341 (2009) | Brain, Human, T2 | no concentration |
| Fleysher et al 2010 | NMR Biomed. 2010 Dec;23(10):1191-8 | Sequence Development (TQF) | Phantom |
| Foy & Burstein 1992 | Magn Reson Med. 1992 Oct;27(2):270-83. | Cardiac, Rat | organ other than brain |
| Gandini Wheeler-Kingshott et al 2018 | Front. Neurosci. 12:810 | Human, Brain, Task-based | Parenchyma/ anatomical attribution not possible |
| Gast et al 2019 | Magn Reson Med. 2019;00:1–9. | B0 shimming | no concentration |
| Gilles et al 2017 | Sci Rep 7: 17435 (2017) | Human, Brain | Parenchyma/ anatomical attribution not possible |
| Gnahm & Nagel 2015 | NeuroImage 105 (2015) 452–461 | Brain, Human, Multiple sclerosis, PVC | no control concentration |
| Gnahm et al 2014 | Magnetic Resonance in Medicine 71:1720–1732 (2014) | Human, reconstruction dev, anatomical constraints | no concentration |
| Goodman et al 2005 | MRM 53:1040–1045 (2005) | Rat (dead) | Parenchyma/ anatomical attribution not possible |

| | | | |
|---|---|---|---|
| Granot et al 1986 | J Magn Reson 1986; 68:575–581 | Brain, Human | no concentration |
| Granot et al 1988 | Radiology. 1988 May;167(2):547-50 | signal intensity | no concentration |
| Grapperon et al 2019 | Radiology 2019; 00:1–9 | Human, Brain, Amyotrophic Lateral Sclerosis | Parenchyma/ anatomical attribution not possible |
| Greiser et al 2005 | J Magn Reson Imaging. 2005 Jan;21(1):78-81. | Cardiac | organ other than brain |
| Grist et al 2018 | Journal of the Neurological Sciences 387 (2018) 111–114 | Brain, Human, Multiple sclerosis | no control concentration |
| Grodd & Klose 1988 | Neuroradiology (1988) 30:399-407 | Brain, Human, Herpes Simplex, Tumour, Stroke | no concentration |
| Grossman et al 1988 | Radiology. 1988 Nov;169(2):305-9. | signal intensity | no concentration |
| Gupta & Gupta 1982 | J. Magn. Reson. 47, 344–350. | erythrocyte preparation, shift reagent dev | cultured tissue / in vitro |
| Ha et al 2018 | Journal of Magnetic Resonance 286 (2018) 110–114 | Coil development, UHF | no concentration |
| Haeger et al 2020 | Alzheimer's Dement. 2020;6:e12032 | Brain, Human, Alzheimer's | Review/non-experimental (protocol, no results) |
| Hancu et al 1999 | Magn Reson Med 42:1146–1154 (1999) | Sequence Development (TQF) | no concentration |
| Hancu et al 2000 | J Magn Reson. 2000 Dec;147(2):179-91 | Cartilage, Bovine | organ other than brain |
| Haneder et al 2012 | Strahlenther Onkol. 2012 Dec;188(12):1146-54. | Kidney | organ other than brain |
| Haneder et al 2015 | Neuroradiology (2015) 57:321–326 | Human, Glioblastoma, case study (no control) | no concentration |
| Harrington et al 2011 | Cephalalgia. 2011 September ; 31(12): 1254–1265 | Rat, Migraine | Parenchyma/ anatomical attribution not possible |
| Hashimoto et al 1991 | Am J Physiol Imaging. 1991;6(2):74-80. | signal intensity | no concentration |
| Heiler et al 2011 | J Magn Reson 34:935–940 (2011) | Mouse, Brain, Stroke | Animal model, no concentration |
| Henzler et al 2012 | Rofo. 2012 Apr;184(4):340-4. doi | Lung, human, cancer | organ other than brain |
| Hilal et al 1983 | AJNR 4:245-249 1983 | Brain, Cat, Stroke | Animal model, no concentration |
| Hilal et al 1985 | J Comput Assist Tomogr. 1985 Jan-Feb;9(1):1-7. | signal intensity | no concentration |
| Hilgenberg & Smith 2007 | J Vis Exp. 2007;(10):562 | in vitro | cultured tissue / in vitro |
| Hillenbrand et al 2005 | Magn Reson Med. 2005 Apr;53(4):843-50. | Cardiac, dog | organ other than brain |
| Hoesl et al 2019 | Magn Reson Med. 2020 84: 2412-2428 | Sequence Development CRISTINA, human volunteers, phantom | no concentration |
| Hoesl et al 2019 | Magn Reson Med . 2020 Nov;84(5):2412-2428. | Sequence development, CRISTINA | no concentration |

| | | | |
|---|---|---|---|
| Hofeler et al 1987 | Biochemistry. 1987 Aug 11;26(16):4953-62. | Yeast | cultured tissue / in vitro |
| Horn 2004 | Curr Vasc Pharmacol 2004;2: 329–33 | Cardiac, Review | Review/non-experimental |
| Horn et al 2001 | Magn Reson Med. 2001 May;45(5):756-64. | Cardiac, Dog, Rat | organ other than brain |
| Huang et al 2018 | Cureus 10(4): e2502 | Human, Brain, Tumour | no control concentration |
| Hubbard 1970 | 1970;53:985–987 | Theory, no concentrations | Review/non-experimental |
| Huhn et al 2017 | Journal of the Neurological Sciences 379 (2017) 163–166 | Brain, Human, Multiple sclerosis | no control concentration |
| Hui et al 2016 | Neurochem Int. 2016 Mar;94:23-31 | in vitro | cultured tissue / in vitro |
| Hussain et al 2009 | Ann Neurol 2009;66:55–62 | Human, Brain, Stroke | no concentration |
| Hutchison & Shapiro 1991 | Concepts in Magnetic Resonance, 1991, 3, 215-236 | Review | Review/non-experimental |
| Hutchison et al 1990 | J Biol Chem. 1990 Sep 15;265(26):15506-10. | Bovine serum, DQF | cultured tissue / in vitro |
| Hutchison et al 1993 | Magn Reson Med. 1993 Mar;29(3):391-5. | Cardiac | organ other than brain |
| Inglese et al 2013 | Multiple Sclerosis and Related Disorders (2013) 2, 263–269 | Review, MS | Review/non-experimental |
| Inglese et al 2018 | Expert Rev Neurother. 2018 Mar;18(3):221-230 | Review, MS | Review/non-experimental |
| Insko et al 1997 | J Magn Reson Imaging. 1997 Nov-Dec;7(6):1056-9. | Cartilage | organ other than brain |
| Insko et al 2002 | Acad Radiol. 2002 Jul;9(7):800-4. | Cartilage/Skeletal, Spine | organ other than brain |
| Jacobs et al 2009 | J Magn Reson Imaging. 2009 Mar;29(3):649-56 | Uterus | organ other than brain |
| Jans et al 1988 | Magn Reson Med. 1988 Jul;7(3):292-9 | renal cell preparation | cultured tissue / in vitro |
| Jansen et al 2004 | Circulation. 2004 Nov 30;110(22):3457-64. | Cardiac | organ other than brain |
| Jelicks & Gupta 1989 | J. Magn. Res. 81, 586–592. | erythrocyte preparation | cultured tissue / in vitro |
| Jerecic et al 2002 | Biomed Tech (Berl). 2002;47 Suppl 1 Pt 1:458-9. | Cardiac | organ other than brain |
| Jerecic et al 2004 | MAGMA. 2004 May;16(6):297-302 | Cardiac | organ other than brain |
| Jones et al 2006 | Stroke. 2006 Mar;37(3):883-8 | Rat, Brain, Stroke | Animal model, no concentration |
| Jospeh & Summers 1987 | MRM 467-77 (1987) | Methodology | no concentration |
| Kaffanke et al 2009 | J Magn Reson. 2009 Aug;199(2):117-25 | Sequence development, SPRITE | Phantom/ Simulation |
| Kahn et al 2021 | Sci Rep. 2021 Mar 23;11(1):6710. | Rat, Brain, Glioma, MRSI | Animal model |
| Kalyanapuram et al 1998 | J Magn Reson Imaging. 1998 Sep-Oct;8(5):1182-9. | Dog, TQF | Animal model, no concentration |
| Kaplan et al 1997 | Cancer Res. 1997 Apr 15;57(8):1452-9. | Pancreas, Various animal models, Cancer | organ other than brain |
| Kauczor & Kreitner 1999 | Eur Radiol. 1999;9(9):1755-64. | Cardiac | organ other than brain |

| | | | |
|---|---|---|---|
| Keltner et al 1994 | J Magn Reson B. 1994 Jul;104(3):219-29. | Sequence Development (TQF) | Phantom |
| Kemp-Harper et al 1995 | J. Magn. Reson. B 108, 280–284. | Sequence development, TQF, phantom study, mussel | Phantom / non-human |
| Kim et al 1997 | Circulation. 1997 Apr 1;95(7):1877-85. | Cardiac, Rabbit | organ other than brain |
| Kim et al 1999 | Circulation. 1999 Jul 13;100(2):185-92. | Cardiac, Rabbit and dog | organ other than brain |
| Kirsch et al 2010 | NMR Biomed. (2010) DOI:10.1002/nbm.1500 | Rat, Multi-nuclear imaging | Animal model, no concentration |
| Kleimaier et 2020 | NMR Biomed. 2020 Oct;33(10):e4367 | Bovine serum, TQF | cultured tissue / in vitro |
| Kleimaier et al 2020 | NMR in Biomedicine. 2020;e4367 | TQF, bovine serum albumin | cultured tissue / in vitro |
| Kleinhans et al 2014 | J Vis Exp. 2014 Oct 8;(92):e52038. | Mouse, multi-photon fluorescence microscopy | non Na-MRI, animal model |
| Kline et al 2000 | Clin Cancer Res. 2000 Jun;6(6):2146-56. | Pancreas, Mouse, cancer | organ other than brain |
| Knubovets et al 1998 | J. Magn. Reson. 131, 92–96 | erythrocyte preparation | cultured tissue / in vitro |
| Koehler et al 2001 | MAGMA. 2001 Oct;13(2):63-9. | Human, Brain | Parenchyma/ anatomical attribution not possible |
| Kohler et al 1989 | J Magn Reson 1989:83:423–427. | No access | No access |
| Kohler et al 1992 | Magn Reson Med. 1992 Jan;23(1):77-88. | Ocular, Rabbit | organ other than brain |
| Kolbe et al 2020 | NeuroImage 211 (2020) 116609 | Brain, Human, T2 | no concentration |
| Kolodny et al 1987 | Arch Ophthalmol. 1987;105(11):1532-1536 | Ocular tissue | organ other than brain |
| Kolodny et al 1989 | Surv Ophthalmol. 1989 May-Jun;33(6):502-14. | Ocular tissue | organ other than brain |
| Kolodny et al 1993 | Invest Ophthalmol Vis Sci. 1993 May;34(6):1917-22. | Ocular, Rabbit, Human | organ other than brain |
| Konstandin & nagel 2014 | MAGMA. 2014 Feb;27(1):1-4. | Review, Imaging fast relaxing Nuclei | Review/non-experimental |
| Konstandin & Nagel 2014 | Magn Reson Mater Phy (2014) 27:5–19 | Review, methodology, UTE imaging | Review/non-experimental |
| Konstandin et al 2011 | Magn Reson Med. 2011 Apr;65(4):1090-6 | Sequence development, DA-2D | no concentration |
| Konstandin et al 2015 | Magnetic Resonance Imaging 33 (2015) 319–327 | Spine, Heart, Simulation, Phantom | organ other than brain |
| Kratzer et al 2021 | Magn Reson Med. 2021;00:1–14. | Human, Brain, Relaxometry | no concentration |
| Lachner et al 2019 | Z Med Phys. 2019 Dec 19. pii: S0939-3889(19)30126-6 | Brain, Human | no concentration |
| Lakshmanan et al 2018 | NMR Biomed. 2018 Feb;31(2) | Coil design | no concentration |
| Langer & Rose 2009 | J Physiol. 2009 Dec 15;587(Pt 24):5859-77 | astrocyte culture study | cultured tissue / in vitro |
| LaVerde et al 2007 | Magn Reson Med 57:201–205 (2007) | Primate, Brain, Stroke | Animal model, no concentration |
| LaVerde et al 2009 | JMRI 30:219–223 (2009) | Monkey, Brain, Stroke | Animal model |

| | | | |
|---|---|---|---|
| Laymon et al 2012 | Magn Reson Imaging. 2012 November ; 30(9) | Human, Glioblastoma | Parenchyma/ anatomical attribution not possible |
| Lee et al 1986 | Magn Reson Imaging. 1986;4(4):343-50 | signal intensity | no concentration |
| Lee et al 2000 | Magn Reson Med. 2000 Feb;43(2):269-77. | Cardiac tissue | organ other than brain |
| Liebling & Gupta 1987 | Ann NY Acad Sci 1987; 508: 149-163 | Human, Neoplastic & Non-neoplastic excised tissue | cultured tissue / in vitro |
| Lim et al 1989 | NMR Biomed. 1989 Sep;2(3):120-3. | 23Na methodology, coil design | no concentration |
| Lin et al 2001 | Stroke 32(4):925-32 2001 | Rat, Brain, Stroke | Animal model, no concentration |
| Lommen et al 2016 | NMR Biomed. 2016; 29: 129–136 | Human, Brain | Parenchyma/ anatomical attribution not possible |
| Lommen et al 2018 | Magnetic Resonance in Medicine 80:571–584 (2018) | Brain, Human, T2 | no concentration |
| Lu et al 2011 | Journal of Magnetic Resonance 213 (2011) 176–181 | Sequence development (flexTPI) | no concentration |
| Lu et al 2019 | Journal of Magnetic Resonance 307 (2019) 106582 | Motion correction | no concentration |
| Lupu et al 2009 | Photodiagnosis Photodyn Ther. 2009 Sep-Dec;6(3-4):214-20 | Retinal tumour | organ other than brain |
| Lyon et al 1991 | MRM 18,80-92 ( 1991 ) | Brain, Rat | Animal model, no concentration |
| Madelin & Regatte et al 2013 | J Magn Reson Imaging. 2013 Sep;38(3):511-29 | Review, Biomedical applications | Review/non-experimental |
| Madelin et al 2014 | Prog. Nucl. Magn. Reson Spectrosc. 79, 14–47. | Review, Methodology | Review/non-experimental |
| Maguire et al 2015 | J Cardiovasc Magn Reson. 2015 Jun 15;17:45 | Cardiac, Mouse | organ other than brain |
| Malliard 2010 | Ann Pharm Fr. 2010 May;68(3):195-202 | Retinoblastoma | organ other than brain |
| Mallow et al 1990 | Magn Reson Med. 1990 Jul;15(1):33-44. | Cardiac, Rat | organ other than brain |
| Malzacher et a 2019 | MRI 59 (2019) 97-104 | Coil development | no concentration |
| Malzacher et al 2016 | Z Med Phys. 2016 Mar;26(1):95-100 | Human, spine | organ other than brain |
| Maril et al 2006 | Kidney Int. 2006 Feb;69(4):765-8. | Kidney, Rat | organ other than brain |
| Matthies et al 2010 | J Magn Reson. 2010 Feb;202(2):239-44 | Sequence Development (TQF) | Phantom |
| Matwiyoff et al 1986 | Magn Reson Med. 1986 Feb;3(1):164-8. | Muscle, rat | organ other than brain |
| Maudsley & Hilal 1984 | British Medical Bulletin (1984) Vol. 40, No. 2, pp. 165-166 | Review | Review/non-experimental |
| Mellon et al 2009 | AJNR Am J Neuroradiol. 2009 May;30(5):978-84 | Brain, Human, Alzheimers | no concentration |
| Mirkes et al 2016 | Magnetic Resonance in Medicine 75:1278–1289 (2016) | Human, Brain, UHF, TQF | no concentration |
| Miyazaki & Ross 2015 | eNeuro. 2015 Nov 9;2(5). pii: ENEURO.0092-15.2015 | Brain, Rat, fluorescence imaging | non Na-MRI, animal model |
| Mohamed et al 2020a | In Vivo . Jan-Feb 2021;35(1):429-435 | Brain, Human, Alzheimer's | no concentration |

| | | | |
|---|---|---|---|
| Mohamed et al 2020b | J Neuroimaging 2020;0:1-9. | Human, Brain, Tumour | no control concentration |
| Mori et al 2000 | NEUROSURGERY (2000) 46:1 157-168 | Rat, Epilepsy | Animal model |
| Moseley et al 1985 | Magn Reson Imaging. 1985;3(4):383-7. | Rat, Brain, Stroke | Animal model, no concentration |
| Moshrefi-Ravasdjani et al 2017 | Neurochem Res. 2017 Sep;42(9):2505-2518 | astrocytes, neurochemical study | cultured tissue / in vitro |
| Nagel et al 2009 | Magn Reson Med. 2009 Dec;62(6):1565-73 | Sequence development, DA-3D-RAD | no concentration |
| Nagel et al 2011 | Invest Radiol 2011;46: 539–547 | Human, Brain, Tumour | Parenchyma/ anatomical attribution not possible |
| Nansky et al 2005 | Neoplasia. 2005 Jul; 7(7): 658–666. | Mouse, Tumour | Parenchyma/ anatomical attribution not possible |
| Naritomi et al 1987 | Biophys J. 1987 Oct;52(4):611-6. | Gerbil, Brain | Animal model, no concentration |
| Naritomi et al 1988 | J Cereb Blood Flow Metab. 1988 Feb;8(1):16-23 | Gerbil | Animal model, no concentration |
| Navon et al 1993 | Magn Reson Med 1993;30:503–506. | Cardiac, Rat | organ other than brain |
| Navon et al 2001 | NMR Biomed. 2001;14:112–132 | Review, MQF | Review/non-experimental |
| Neuberger et al 2004 | MAGMA. 2004 Dec;17(3-6):196-200. | Cardiac, Mouse | organ other than brain |
| Neumaier-Probst et al 2015 | Int J Stroke. 2015 Oct;10 Suppl A100:56-61 | Human, Brain, Stroke | no control concentration |
| Nielles-Vallespin et al 2007 | Magn Reson Med 57:74–81 (2007) | Human, Glioma | no concentration |
| Niesporak et al 2017 | MAGMA. 2017 Dec;30(6):519-536 | Partial Volume Correction , T2 | no concentration |
| Noebauer-Huhmann et al 2012 | Radiology. 2012 Nov;265(2):555-64 | Human, spine | organ other than brain |
| Nunes Neto et al 2018 | Neuroradiology. 2018 August ; 60(8): 795–802 | Human, Glioma | no controls |
| Onizuka et al 2004 | Anesthesiology. 2004 Jul;101(1):110-20. | patch clamp | cultured tissue / in vitro |
| Onizuka et al 2011 | Anesth Analg. 2011 Mar;112(3):703-9 | patch clamp | cultured tissue / in vitro |
| Ouwerkerk & Morgan 2007 | J Am Coll Radiol. 2007;4:739. | Review | Review/non-experimental |
| Ouwerkerk 2011 | Methods Mol Biol. 2011;711:175-201 | Review | Review/non-experimental |
| Ouwerkerk et al 2007 | Breast Cancer Res Treat. 2007 Dec;106(2):151-60 | Breast | organ other than brain |
| Parish et al 1997 | Magn Reson Med. 1997 Oct;38(4):653-61. | Cardiac, Human | organ other than brain |
| Paschke et al 2018 | Invest Radiol. 2018 Sep;53(9):555-562 | Human, Brain, Stroke | no control concentration |
| Pathak et al 1995 | Am J Surg. 1995 Nov;170(5):423-6. | Cheek Tissue, Hamster, fluorescence imaging | organ other than brain |

| | | | |
|---|---|---|---|
| Pekar & Leigh 1986 | J. Magn. Reson. 69, 582–584 | DQF, bovine serum albumin | cultured tissue / in vitro |
| Pekar et al 1987 | J Magn Reson 1987; 72 (1): 159-161 | dog, erythrocyte preparation, DQF | cultured tissue / in vitro |
| Perman et al 1986 | Radiology. 1986 Sep;160(3):811-20. | signal intensity /relaxation (monoexp) | no concentration |
| Perman et al 1989 | Magn Reson Med. 1989 Feb;9(2):153-60 | Sequence development (MultiTE) | no concentration |
| Petracca et al 2016 | NMR Biomed. 2016 Feb;29(2):153-61 | Review, MS | Review/non-experimental |
| Pike et al 1984 | Am J Physiol. 1984 May;246(5 Pt 1):C528-36. | erythrocyte preparation | cultured tissue / in vitro |
| Pinker et al 2011 | Breast Care (Basel). 2011;6(2):110-119 | Review, breast, cancer | Review/non-experimental |
| Pinker et al 2012 | Eur J Radiol. 2012 Mar;81(3):566-77 | Review, cancer | Review/non-experimental |
| Qian et al 2009 | Magnetic Resonance Imaging 27 (2009) 656–663 | Sequence development (parallel imaging TPI) | no concentration |
| Qian et al 2010 | Magn Reson Med. 63:543–552 | Sequence Development (UHF) | no concentration |
| Qian et al 2012 | Magn Reson Med. 68:1808–1814 (2012) | Brain, Human, UHF | no concentration |
| Qian et al 2015 | Magn Reson Med. 2015 Jul;74(1):162-174 | Human, Brain, Lesion | Parenchyma/ anatomical attribution not possible |
| Ra et al 1986 | Magn Reson Med. 1986 Apr;3(2):296-302. | Sequence development (UTE) | no concentration |
| Ra et al 1988 | MRM 7, 11-22 (1988) | Human, heart, liver, gallbladder, kidney, and spine | organ other than brain |
| Ra et al 1989 | J Comput Assist Tomogr. 1989 Mar-Apr;13(2):302-9. | Methodology | no concentration |
| Reddy et al 1995 | MRM 33.134-139 (1995) | Human, Brain, Skeletal Muscle | no concentration |
| Reddy et al 1997 | Magn Reson Med. 1997 Aug;38(2):279-84. | Cartilage, bovine | organ other than brain |
| Reddy et al 1998 | Magn Reson Med. 1998 May;39(5):697-701. | Cartilage, bovine | organ other than brain |
| Reimer et al 2014 | Magn Reson Mater Phy (2014) 27:35–46 | Sequence development (3D cones) | no concentration |
| Reimer et al 2018 | NMR Biomed. 2018 May;31(5) | Brain, Human, T2 | no concentration |
| Remele et al 2017 | Cereb Cortex. 2017 Jun 1;27(6):3272-3283 | patch clamp, astrocytes | cultured tissue / in vitro |
| Ridley et al 2017 | NeuroImage 157 (2017) 173–183 | Human, Brain, Epilepsy | Parenchyma/ anatomical attribution not possible |
| Romanzetti et al 2006 | Journal of Magnetic Resonance 179 (2006) 64–72 | Sequence development (SPRITE) | no concentration |
| Romanzetti et al 2014 | Neuroimage. 96, 44–53. | Human, Brain, UHF sequence comparison | no concentration |
| Ronen & Kim 2001 | NMR Biomed. 2001 Nov-Dec;14(7-8):448-52. | Rat, shift reagent | Animal model |
| Rong et al 2008 | J Magn Reson. 2008 Aug;193(2):207-9 | Cartilage | organ other than brain |

| | | | |
|---|---|---|---|
| Rooney & Springer 1991 | NMR IN BIOMEDICINE, VOL. 4,227-245 (1991) | bovine serum, yeast | cultured tissue / in vitro |
| Rooney & Springer 1991 | NMR Biomed. 4 (1991) 209–226 | Review | Review/non-experimental |
| Rose & Verkhratsky 2016 | Glia. 2016 Oct;64(10):1611-27 | Review, astrocytes | Review/non-experimental |
| Rose 2012 | Cold Spring Harb Protoc. 2012 Nov 1;2012(11):1161-5 | two-photon imaging | non Na-MRI |
| Ross et al 1993 | Jpn J Physiol. 1993;43 Suppl 1:S83-9. | electrophysiology | cultured tissue / in vitro |
| Ross et al 2000 | Ann N Y Acad Sci. 2000 May;904:12-7. | Review, muscle imaging | Review/non-experimental |
| Sakuta et al 2020 | Pflugers Arch . 2020 May;472(5):609-624 | Knockout mice | cultured tissue / in vitro |
| Sandstede et al 2000 | Rofo. 2000 Sep;172(9):739-43. | Cardiac | organ other than brain |
| Sandstede et al 2004 | Magn Reson Med. 2004 Sep;52(3):545-51. | Cardiac | organ other than brain |
| Scelfo et al 2003 | J Neurophysiol. 2003 May;89(5):2555-63. | electrophysiology | cultured tissue / in vitro |
| Schepkin et al 1996 | J Appl Physiol (1985). 1996 Dec;81(6):2696-702. | Cardiac, Rat | organ other than brain |
| Schepkin et al 1998 | Magn Reson Med. 1998 Apr;39(4):557-63. | Cardiac, Rat | organ other than brain |
| Schepkin et al 2005 | MRM 53:85–92 | Rat, Glioma | Parenchyma/ anatomical attribution not possible |
| Schepkin et al 2006 | Magn Reson Imaging. 2006 Apr;24(3):273-8 | Rat, Subcutaneous tumour | Animal model |
| Schepkin et al 2006 | NMR Biomed. 2006 December ; 19(8): 1035–1042 | Rat, Subcutaneous tumour | Animal model |
| Schepkin et al 2010 | Magn Reson Imaging. 2010 April ; 28(3): 400–407 | Rat, UHF | Animal model, no concentration |
| Schepkin et al 2012 | Magn Reson Med. 67:1159–1166 (2012) | Rat, Glioma | Animal model |
| Schepkin et al 2013 | Magn Reson Mater Phy 27(1) June 2013 | Rat, Brain | Parenchyma/ anatomical attribution not possible |
| Schepkin et al 2016 | NMR Biomed. 2016 Feb;29(2):175-86 | Review, Glioma, Animal models | Review/non-experimental |
| Schnall et al 1988 | Magn Reson Med. 6, 15-23 (1988) | Cat, NMR spectroscopy, Epilepsy | Animal model, no concentration |
| Schuierer et al 1991 | Magn Reson Med. 1991 Nov;22(1):1-9. | signal intensity | no concentration |
| Seshan et al 1995 | Magn Reson Med. 1995 Jul;34(1):25-31. | Kidney, Rat | organ other than brain |
| Seshan et al 1997 | Magn Reson Med. 1997 Nov;38(5):821-7. | Liver, Rat | organ other than brain |
| Shah et al 2009 | Journal of Magnetic Resonance 199 (2009) 136–145 | SPRITE, phantom & PSF simulation | no concentration |
| Shah et al 2016 | NMR Biomed. 2016 Feb;29(2):162-74. | Review, Brain | Review/non-experimental |
| Shajan et al 2016 | Magnetic Resonance in Medicine 75:906–916 (2016) | Human, Brain, UHF, Coil design, | no concentration |
| Shapiro et al 2002 | Magn Reson Med. 2002 Feb;47(2):284-91. | Cartilage | organ other than brain |

| | | | |
|---|---|---|---|
| Shen et al 1997 | Magn Reson Med. 1997 Nov;38(5):717-25. | 23Na methodology, coil design | no concentration |
| Shimizu et al 1992 | Neuroradiology. 1992;34(4):301-4. | Human, Brain, Stroke | no concentration |
| Shimizu et al 1993 | Neuroradiology. 1993;35(6):416-9. | Human, Brain, Stroke | no concentration |
| Shinar & Navon 1984 | Biophys Chem. 1984 Nov;20(4):275-83. | erythrocyte preparation | cultured tissue / in vitro |
| Shymanskaya et al 2019 | Mol Imaging Biol (2019) | Human, Glioma | no control concentration |
| Smith et al 1990 | Int Ophthalmol. 1990 Mar;14(2):119-24. | Ocular applications | organ other than brain |
| Solanky et al 2013 | Magn Reson Med2013;69:1201–8. | Spine | organ other than brain |
| Springer 1987 | Annu Rev Biophys Biophys Chem. 1987;16:375-99 | review | Review/non-experimental |
| Staroswiecki et al 2010 | J Magn Reson Imaging. 2010 Aug;32(2):446-51. | Cartilage | organ other than brain |
| Steidle et al 2004 | Magn Reson Imaging. 2004 Feb;22(2):171-80. | Torso | organ other than brain |
| Steiner 1986 | Magn Reson Med. 1986 Aug;3(4):473-90. | review | Review/non-experimental |
| Stelzeneder & Trattnig 2010 | Radiologe. 2010 Dec;50(12):1115-9 | Human, spine | organ other than brain |
| Stobbe & Beailieu 2018 | Magnetic Resonance in Medicine 79:2968–2977 (2018) | Brain, Human, Multiple sclerosis | no control concentration |
| Stobbe & Beaulieu 2005 | Magn Reson Med 54:1305–1310 (2005) | Sequence development, IR | no concentration |
| Stobbe & Beaulieu 2008 | Magn Reson Imaging. 59:345–355 (2008) | Sequence design, high field | no concentration |
| Stobbe & Beaulieu 2008 | Magn Reson Imaging. 60:981–986 (2008) | Sequence development (3D-TPI, DA) | no concentration |
| Stobbe & Beaulieu 2014 | Magn Reson Mater Phy (2014) 27:21–33 | Sequence development (PACMAN) | no concentration |
| Stobbe & Beaulieu 2016 | NMR Biomed. 2016 Feb;29(2):119-28 | Methodology, residual quadrupole effects | no concentration |
| Summers et a 1991 | Invest Radiol 1991 26: 233-241 | Brain, Mouse, Neuroblastoma | Animal model, no concentration |
| Summers et al 1988 | MRM 8,427-439 (1988) | Rat | organ other than brain |
| Syeda et al 2019 | Magn Reson Med. 2019;1–11. | Human, Brain, Relaxometry | no concentration |
| Szmacinski & Lakowicz 1997 | Anal Biochem. 1997 Aug 1;250(2):131-8. | fluorescence imaging | non Na-MRI |
| Tannous et al 2009 | Mol Ther. 2009 May;17(5):810-9. | Sodium channel, culture study | cultured tissue / in vitro |
| Tauskela et al 1997 | J Magn Reson. 1997 Jul;127(1):115-27 | Cardiac, Rat | organ other than brain |
| Tenase & Boada 2005 | Journal of Magnetic Resonance 174 (2005) 270–278 | Sequence development, TQF, phantom study | Phantom |
| Thulborn & Ackerman 1983 | J. Magn. Reson. 55, 357–371 | NMR, Molar concentration of metabolites | no concentration |
| Thulborn et al 1997 | Int J Imaging Syst Technol, **8,** 572–581, 1997 | Human, Brain, Stroke, Epilepsy | no concentration |
| Thulborn et al 1998 | Magn Reson Med. 1998 Mar;39(3):369-75. | Methodology, B1 inhomogeneities | no concentration |

| | | | |
|---|---|---|---|
| Thulborn et al 1999 | Radiology 1999;139:26– 34 | Brain, Human, Primate, Glioma | no controls |
| Thulborn et al 1999 | Radiology. 1999 Oct;213(1):156-66. | Human, Brain, Stroke | Parenchyma/ anatomical attribution not possible |
| Thulborn et al 1999 | MRM 41:351–359 (1999) | Rat, Glioma | Parenchyma/ anatomical attribution not possible |
| Thulborn et al 2002 | Comput Med Imaging Graph 2001;16:73– 89 | image analysis software | Review/non-experimental |
| Thulborn et al 2009 | Neuroimaging Clin N Am. 2009 ; 19(4): 615–624 | Review, Brain Tumours | Review/non-experimental |
| Thulborn et al 2018 | NeuroImage 168 (2018) 250–268 | Review, Biomedical applications | Review/non-experimental |
| Thulborn et al 2019 | Clin Cancer Res. 2019 Feb 15;25(4):1226-1232 | Human, Glioma | no controls |
| Tofts & Wray 1988 | NMR Biomed. 1, 1–10. | Review | Review/non-experimental |
| Trattnig 1997 | Eur J Radiol. 1997 Nov;25(3):188-98. | Cartilage | organ other than brain |
| Trattnig et al 2006 | Z Rheumatol. 2006 Dec;65(8):681-7. | Review, Joint imaging | Review/non-experimental |
| Trattnig et al 2010 | Eur Radiol. 2010 Aug;20(8):2039-46 | Knee joint, human | organ other than brain |
| Trattnig et al 2012 | Eur Radiol. 2012 Nov;22(11):2338-46 | Cartilage, Human | organ other than brain |
| Trattnig et al 2016 | NMR Biomed. 2016 Sep;29(9):1316-34 | Review, UHF | Review/non-experimental |
| Trattnig et al 2019 | Radiologe. 2019 Aug;59(8):742-749 | Cartilage, Review, German | Review/non-experimental |
| Truong et al 2014 | J Magn Reson. 2014 October ; 247: 88–95 | Rat, Migraine | Animal model |
| Tsang et al 2011 | JMRI 33:41–47 (2011) | Human, Brain, Stroke | no concentration |
| Tsang et al 2012 | Magnetic Resonance in Medicine 67:1633–1643 (2012) | Sequence development (TQF) | no concentration |
| Tsang et al 2013 | Journal of Magnetic Resonance 230 (2013) 134–144 | Sequence Development (TQF) | no concentration |
| Tsang et al 2015 | Magn Reson Med. 2015 Feb;73(2):497-504 | Sequence development (DQF) | no concentration |
| Turski et al 1986 | Radiology 1986; 160:821-825 | Dog, Oedema | Animal model, no concentration |
| Turski et al 1987 | Radiology. 1987 Apr;163(1):245-9. | Dog, Human, Neoplasm | Animal model, no concentration |
| Turski et al 1988 | Radiol Clin North Am. 1988 Jul;26(4):861-71. | No access | No access |
| Tyson et al 1996 | Stroke. 1996;27:957–964 | Rat, Brain, Stroke | Animal model, no concentration |
| van der Veen et al 1993 | Magn Reson Med. 1993 Apr;29(4):571-4. | erythrocyte preparation | cultured tissue / in vitro |
| Veliyulin et al 2009 | J Agric Food Chem. 2009 May 27;57(10):4091-5 | Food study | organ other than brain |
| Vinitski et al 1987 | MRM 5, 278-285 (1987) | signal intensity /relaxation (monoexp) | no concentration |
| Virapongse et al 1989 | Top Magn Reson Imaging. 1989 Dec;2(1):63-75. | Review, Stroke, MRI applications | Review/non-experimental |

| | | | |
|---|---|---|---|
| Wang et al 1996 | Epilepsia, 37( 10):1000-1006, 1996 | Rat, Epilepsy | Animal model |
| Wang et al 2000 | Stroke. 2000 Jun;31(6):1386-91; | Rat, Brain, Stroke | Animal model |
| Wang et al 2009 | J Magn Reson Imaging. 2009 Sep;30(3):606-14 | Knee joint, human | organ other than brain |
| Wang et al 2010 | Spine 2010;35:505–10. | Spine | organ other than brain |
| Weber et al 2010 | Invest Radiol. 2010 Dec;45(12):755-68 | Brain, Human, Glioma | no controls |
| Weber et al 2011 | Neurology. 2011 Dec 6;77(23):2017-24 | Muscle, Human, Duchenne Muscular Dystrophy | organ other than brain |
| Weinberg et al 1991 | Invest Ophthalmol Vis Sci. 1991 Jul;32(8):2212-8. | Ocular, Rabbit | organ other than brain |
| Weingärtner et a 2015 | Z. Med.Phys. 25 (2015) 275–286 | Methodology, scan time reduction | no concentration |
| Wetterling 2015 | Journal of Cerebral Blood Flow & Metabolism (2015) 35, 103–110 | Human, Brain, Stroke | no concentration |
| Wetterling et al 2010 | Physics in Medicine and Biology 55(24):7681-95 | Rat, Brain, Stroke | Animal model |
| Wetterling et al 2012 | Journal of Magnetic Resonance 217 (2012) 10–18 | Coil development | no concentration |
| Wetterling et al 2012 | Phys Med Biol. 2012 Jul 21;57(14):4555-67 | Human, Whole body imaging | organ other than brain |
| Wetterling et al 2012 | Magn Reson Med. 67:740–749 (2012) | Rat, Brain, Stroke | Animal model |
| Wetterling et al 2016 | BMC Neurosci (2016) 17:82 | Rat, Brain, Stroke | Animal model, no concentration |
| Wiggins et al 2016 | NMR Biomed. 2016 Feb;29(2):96-106 | Review, Coil design | Review/non-experimental |
| Wilferth et al 2019 | Magn Reson Imaging. 2019 Nov;63:280-290 | Muscle, Human | organ other than brain |
| Wimperis & Wood 1991 | J. Magn. Res. 95, 428–436 | Sequence Development (TQF) | Phantom |
| Wimperis et al 1992 | J Magn Reson 1992;98:628–636. | Sequence Development (TQF) | Phantom |
| Winkler 1990 | Neuroradiology. 1990;32(5):416-20. | review | Review/non-experimental |
| Winter & Bansal 2001 | MRM: 45 436-442 (2001) | Rat, Tumour concentrations | Animal model, no concentration |
| Winter & Bansal 2001 | J Magn Reson. 2001 Sep;152(1):70-8. | Tumour, Mice, TQF | organ other than brain |
| Winter et al 1998 | J. Appl. Physiol. 85(5): 1806–1812, 1998 | Liver, Rat, Shift Reagent | organ other than brain |
| Winter et al 2001 | CANCER RESEARCH 61, 2002–2007, March 1, 2001] | Rat, Glioma | Animal model, no concentration |
| Worthoff et al 2020 | NMR in Biomedicine. 2020;e4361. https://doi.org/10.1002/nbm.4361 | Brain, Human, Glioma, TQ/SQ, T2 | no control concentration |
| Young et al 1986 | Cent Nerv Syst Trauma. 1986 Summer;3(3):215-34 | Rat, Brain, Stroke, atomic absorption spectroscopy | non Na-MRI, animal model |
| Young et al 1987 | Stroke. 1987 Jul-Aug;18(4):751-9 | Rat,Brain, Stroke, atomic absorption spectroscopy | Animal model, no concentration |
| Yushmanov et al 2007 | Magn Reson Med. 57:494–500 (2007) | Rat, Brain, Stroke | Animal model |

| | | | |
|---|---|---|---|
| Yushmanov et al 2009 | J Magn Reson Imaging 29:962–966 (2009) | Rat, Brain, Stroke | Animal model |
| Yushmanov et al 2009 | J Magn Reson Imaging. 30:18–24 (2009) | Rat, Brain, Stroke | Animal model, no concentration |
| Yushmanov et al 2013 | Brain Research 1527 (2013) 199–208 | Rat, Brain, Stroke | Animal model |
| Zaric et al 2020 | J. MAGN. RESON. IMAGING 2020 DOI: 10.1002/jmri.27326 | Review, applications | Review/non-experimental |
| Zbýň et al 2012 | Osteoarthritis Cartilage. 2012 Aug;20(8):837-45 | Cartilage, Human | organ other than brain |
| Zhang et al 2010 | J Magn Reson. 2010 Jul;205(1):28-37 | Yeast cell suspension | cultured tissue / in vitro |
| Zhao et al 2021 | Magn Reson Med. 2021;86:625–636. | Human, Tumour, Controls | Parenchyma/ anatomical attribution not possible |
| Zuo et al 2006 | J Magn Reson Imaging. 2006 Jul;24(1):191-6. | Muscle | organ other than brain |
| Zuo et al 2008 | Magn Reson Imaging. 2008 Jun;26(5):629-37 | Muscle, Human | organ other than brain |

#### Supplementary Table 4 – COBIDAS domains and coded results indicting if reports were identified (Y=yes, N=no) in included studies.

| Paper | N. | Age | Sex | Handedness | Ethical Approval | Informed Consent | Scanner | Coil | Pulse Sequence | Echo Time (TE) | Repetition Time | Flip Angle (FA) | Acquisition Time | FOV | Nominal Resolution | Software | Basis of segmentation / ROIs | Details of Registration /Normalisation | Information on Smoothing/PSF |
|---|---|---|---|---|---|---|---|---|---|---|---|---|---|---|---|---|---|---|---|
| **Winkler et al 1989** | Y | Y | N | N | Y | Y | Y | N | Y | Y | Y | N | Y | Y | Y | N | N | N | N |
| **Ouwerkerk et al 2003** | Y | Y | Y | N | Y | Y | Y | Y | Y | Y | Y | Y | Y | Y | Y | N | N | Y | N |
| **Thulborn et al 2005** | Y | N | N | N | Y | Y | Y | Y | Y | Y | Y | Y | N | N | Y | Y | N | N | N |
| **Inglese et al 2010** | Y | Y | Y | N | Y | Y | Y | Y | Y | Y | Y | Y | Y | Y | Y | Y | Y | N | N |
| **Lu et al 2010** | Y | Y | Y | N | Y | Y | Y | Y | Y | Y | Y | Y | Y | Y | Y | N | N | N | Y |
| **Reetz et al 2012** | Y | Y | Y | N | Y | Y | Y | Y | Y | Y | Y | Y | Y | Y | Y | Y | Y | Y | N |
| **Qian et al 2012** | Y | Y | Y | N | Y | Y | Y | Y | Y | Y | Y | Y | Y | Y | Y | N | N | N | N |
| **Zaaraoui et 2012** | Y | Y | Y | N | Y | Y | Y | Y | Y | Y | Y | Y | Y | Y | Y | Y | Y | Y | Y |
| **Paling et al 2013** | Y | Y | Y | N | Y | Y | Y | Y | Y | Y | Y | N | Y | Y | Y | Y | Y | Y | N |
| **Maarouf et al 2014** | Y | Y | Y | N | Y | Y | Y | Y | Y | Y | Y | N | Y | Y | Y | Y | Y | Y | Y |
| **Mirkes et al 2015** | Y | Y | Y | N | N | Y | Y | Y | Y | Y | Y | Y | Y | Y | Y | N | N | N | Y |
| **Niesporak et al 2015** | Y | Y | Y | N | Y | Y | Y | Y | Y | Y | Y | Y | Y | Y | Y | Y | Y | Y | N |
| **Eisele et al 2016** | Y | Y | Y | N | Y | Y | Y | Y | Y | Y | Y | Y | Y | N | Y | Y | Y | Y | N |
| **Petracca et al 2016** | Y | Y | Y | N | Y | Y | Y | Y | Y | Y | Y | Y | Y | N | Y | Y | Y | Y | Y |
| **Thulborn et al 2016** | Y | Y | N | N | Y | Y | Y | Y | Y | Y | Y | Y | Y | N | Y | Y | N | N | N |
| **Maarouf et al 2017** | Y | Y | Y | N | Y | Y | Y | Y | Y | Y | Y | N | Y | N | Y | Y | Y | Y | Y |
| **Eisele et al 2017** | Y | Y | Y | N | Y | Y | Y | N | Y | Y | Y | Y | Y | N | Y | N | Y | N | N |
| **Ridley et al 2018** | Y | Y | Y | N | Y | Y | Y | Y | Y | Y | Y | N | Y | N | Y | Y | Y | Y | N |
| **Worthoff et al 2018** | Y | Y | Y | N | Y | Y | Y | Y | Y | Y | Y | Y | Y | Y | Y | Y | Y | N | Y |
| **Reimer et al 2019** | Y | Y | Y | N | Y | Y | Y | Y | Y | Y | Y | N | N | Y | Y | Y | Y | Y | N |
| **Driver et al 2019** | Y | Y | Y | N | Y | Y | Y | Y | Y | Y | Y | Y | Y | N | Y | Y | Y | Y | N |

| | | | | | | | | | | | | | | | | | | | |
|---|---|---|---|---|---|---|---|---|---|---|---|---|---|---|---|---|---|---|---|
| **Liao et al 2019** | Y | Y | Y | N | Y | Y | Y | Y | Y | Y | Y | Y | Y | Y | Y | Y | Y | Y | Y |
| **Meyer et al 2019a** | Y | Y | Y | N | Y | Y | Y | Y | Y | Y | Y | Y | N | Y | Y | Y | Y | Y | N |
| **Meyer et al 2019b** | Y | Y | Y | N | Y | Y | Y | Y | Y | Y | Y | Y | N | N | Y | Y | Y | Y | N |
| **Kim et al 2020** | Y | Y | Y | N | Y | Y | Y | Y | Y | Y | Y | Y | Y | Y | Y | Y | Y | Y | Y |
| **Gerhalter et al 2021** | Y | Y | Y | N | Y | Y | Y | Y | Y | Y | Y | Y | Y | N | Y | Y | Y | Y | Y |
| **Brownlee et al 2019** | Y | Y | Y | N | Y | Y | Y | Y | Y | Y | Y | N | N | N | Y | Y | Y | Y | Y |
| **Schneider et al 2021** | Y | Y | Y | N | Y | Y | Y | Y | Y | Y | Y | Y | Y | N | Y | Y | Y | Y | N |

#### Supplementary Table 5 – Original and recoded Tissue labels

| **Paper** | **Tissue As labelled** | **Tissue as re-coded** |
|---|---|---|
| Thulborn et al 2005 | Dentate Nucleus, Right | WM, Cb+DN |
| Thulborn et al 2005 | Dentate Nucleus, Left | WM, Cb+DN |
| Inglese et al 2010 | WM, Cerebellum | WM, Cb+DN |
| Reetz et al 2012 | Cortex, Cerebellum | WM, Cb+DN |
| Eisele et al 2017 | Dentate Nucleus | WM, Cb+DN |
| Ridley et al 2018 | WM, Cerebellum | WM, Cb+DN |
| Meyer et al 2019a | WM, Cerebellum, pedunculi cerebelli medius | WM, Cb+DN |
| Meyer et al 2019b | WM, Cerebellum, pedunculi cerebelli medius | WM, Cb+DN |
| Ouwerkerk et al 2003 | WM | WM |
| Thulborn et al 2005 | WM average, Right | WM |
| Thulborn et al 2005 | WM average, Left | WM |
| Inglese et al 2010 | WM | WM |
| Lu et al 2010 | WM | WM |
| Qian et al 2012 | WM | WM |
| Zaaraoui et 2012 | WM | WM |
| Paling et al 2013 | WM | WM |
| Maarouf et al 2014 | WM | WM |
| Niesporak et al 2015 | WM | WM |
| Eisele et al 2016 | WM | WM |
| Petracca et al 2016 | WM | WM |
| Maarouf et al 2017 | WM | WM |
| Ridley et al 2018 | WM | WM |
| Worthoff et al 2018 | WM | WM |
| Riemer et al 2019 | WM | WM |
| Driver et al 2019 | WM | WM |
| Liao et al 2019 | WM | WM |
| Meyer et al 2019a | WM | WM |
| Kim et al 2020 | WM | WM |
| Gerhalter et al 2021 | WM | WM |
| Brownlee et al 2019 | WM | WM |
| Thulborn et al 2005 | Thalamus, Right | Thalamus |

| | | |
|---|---|---|
| Thulborn et al 2005 | Thalamus, Left | Thalamus |
| Reetz et al 2012 | Thalamus | Thalamus |
| Thulborn et al 2016 | Thalamus | Thalamus |
| Ridley et al 2018 | Thalamus | Thalamus |
| Liao et al 2019 | Thalamus | Thalamus |
| Gerhalter et al 2021 | Thalamus | Thalamus |
| Schneider et al 2021 | Lateral Geniculate body thal) | Thalamus |
| Schneider et al 2021 | Medial Geniculate body thal) | Thalamus |
| Schneider et al 2021 | Pulvinar thal) | Thalamus |
| Thulborn et al 2005 | Putamen, Right | Putamen |
| Thulborn et al 2005 | Putamen, Left | Putamen |
| Reetz et al 2012 | Putamen | Putamen |
| Ridley et al 2018 | Putamen | Putamen |
| Liao et al 2019 | Putamen | Putamen |
| Gerhalter et al 2021 | Putamen | Putamen |
| Schneider et al 2021 | Putamen | Putamen |
| Thulborn et al 2005 | Cortex, Temporal, Right | GM,Temporal |
| Thulborn et al 2005 | Cortex, Temporal, Left | GM,Temporal |
| Thulborn et al 2005 | Hippocampus, Right | GM,Temporal |
| Thulborn et al 2005 | Hippocampus, Left | GM,Temporal |
| Inglese et al 2010 | GM, Temporal | GM,Temporal |
| Reetz et al 2012 | Amygdala | GM,Temporal |
| Reetz et al 2012 | Hippocampus | GM,Temporal |
| Reetz et al 2012 | Cortex,Temporal | GM,Temporal |
| Liao et al 2019 | GM,Temporal | GM,Temporal |
| Liao et al 2019 | Hippocampus | GM,Temporal |
| Thulborn et al 2005 | Cortex, Parietal, Right | GM, Parietal |
| Thulborn et al 2005 | Cortex, Parietal, Left | GM, Parietal |
| Inglese et al 2010 | GM, Parietal | GM, Parietal |
| Reetz et al 2012 | Precunues | GM, Parietal |
| Reetz et al 2012 | Cortex, Parietal | GM, Parietal |
| Liao et al 2019 | GM, Parietal | GM, Parietal |
| Meyer et al 2019b | GM, postcentral gyri | GM, Parietal |
| Thulborn et al 2005 | Cortex, Occipital, Right | GM, Occipital |
| Thulborn et al 2005 | Cortex, Occipital, Left | GM, Occipital |
| Inglese et al 2010 | GM, Occipital | GM, Occipital |
| Reetz et al 2012 | Cortex, Occipital | GM, Occipital |
| Liao et al 2019 | GM, Occipital | GM, Occipital |
| Thulborn et al 2005 | Cortex, Frontal, Right | GM, Frontal |
| Thulborn et al 2005 | Cortex, Frontal, Left | GM, Frontal |
| Inglese et al 2010 | GM, Frontal | GM, Frontal |
| Reetz et al 2012 | Cortex, Frontal | GM, Frontal |
| Mirkes et al 2015 | GM, prefrontal | GM, Frontal |
| Liao et al 2019 | GM, Frontal | GM, Frontal |
| Winkler et al 1989 | Cortex | GM |
| Ouwerkerk et al 2003 | GM | GM |
| Thulborn et al 2005 | GM average, Right | GM |

| | | |
|---|---|---|
| Thulborn et al 2005 | GM average, Left | GM |
| Inglese et al 2010 | GM, Whole | GM |
| Lu et al 2010 | GM | GM |
| Qian et al 2012 | GM | GM |
| Zaaraoui et 2012 | GM | GM |
| Paling et al 2013 | GM, Cortical | GM |
| Maarouf et al 2014 | GM | GM |
| Niesporak et al 2015 | GM | GM |
| Eisele et al 2016 | GM | GM |
| Petracca et al 2016 | GM | GM |
| Maarouf et al 2017 | GM | GM |
| Ridley et al 2018 | GM | GM |
| Riemer et al 2019 | GM | GM |
| Driver et al 2019 | GM | GM |
| Liao et al 2019 | GM | GM |
| Meyer et al 2019a | GM | GM |
| Kim et al 2020 | GM | GM |
| Gerhalter et al 2021 | GM, Cortical | GM |
| Brownlee et al 2019 | Cortical Grey Matter | GM |
| Reetz et al 2012 | Pallidum | Globus Pallidus |
| Eisele et al 2017 | Globus Pallidus | Globus Pallidus |
| Ridley et al 2018 | Pallidum/Globus Pallidus | Globus Pallidus |
| Gerhalter et al 2021 | Pallidus | Globus Pallidus |
| Schneider et al 2021 | Globus Pallidus | Globus Pallidus |
| Schneider et al 2021 | Ventral Pallidum | Globus Pallidus |
| Winkler et al 1989 | WM, Centrum Semiovale | Deep WM |
| Thulborn et al 2005 | WM,Corona radiata, Right | Deep WM |
| Thulborn et al 2005 | WM,Corona radiata, Left | Deep WM |
| Thulborn et al 2005 | WM,Centrum Semiovale, Right | Deep WM |
| Thulborn et al 2005 | WM, Centrum Semiovale, Left | Deep WM |
| Ridley et al 2018 | WM, Centrum Semiovale | Deep WM |
| Meyer et al 2019b | WM, Centrum Semiovale | Deep WM |
| Gerhalter et al 2021 | Corona radiata | Deep WM |
| Thulborn et al 2005 | WM,Forceps major, Right | Central WM |
| Thulborn et al 2005 | WM,Forceps major, Left | Central WM |
| Thulborn et al 2005 | WM,Forceps minor, Right | Central WM |
| Thulborn et al 2005 | WM,Forceps minor, Left | Central WM |
| Inglese et al 2010 | WM, CC, Splenium | Central WM |
| Inglese et al 2010 | WM, Periventricular | Central WM |
| Ridley et al 2018 | WM, CC | Central WM |
| Gerhalter et al 2021 | CC body | Central WM |
| Gerhalter et al 2021 | CC genu | Central WM |
| Gerhalter et al 2021 | CC splenium | Central WM |
| Reetz et al 2012 | Caudate | Caudate |
| Ridley et al 2018 | Caudate | Caudate |
| Liao et al 2019 | Caudate | Caudate |
| Meyer et al 2019a | Caudate Nucleus, Head | Caudate |

| | | |
|---|---|---|
| Gerhalter et al 2021 | Caudate | Caudate |
| Schneider et al 2021 | Caudate | Caudate |
| Thulborn et al 2005 | Midbrain, Right | Brainstem+Pons |
| Thulborn et al 2005 | Midbrain, Left | Brainstem+Pons |
| Thulborn et al 2005 | Brainstem, Right | Brainstem+Pons |
| Thulborn et al 2005 | Brainstem, Left | Brainstem+Pons |
| Thulborn et al 2016 | WM, Brainstem | Brainstem+Pons |
| Ridley et al 2018 | WM, Brainstem, Pons | Brainstem+Pons |
| Meyer et al 2019a | WM, Brainstem, Pons | Brainstem+Pons |
| Meyer et al 2019b | WM, Brainstem, Pons | Brainstem+Pons |
| Schneider et al 2021 | Red Nucleus | Brainstem+Pons |
| Schneider et al 2021 | Locus Coeruleus | Brainstem+Pons |
| Schneider et al 2021 | Ventral Tegmental area | Brainstem+Pons |
| Schneider et al 2021 | Dorsal raphe nucleus | Brainstem+Pons |
| Schneider et al 2021 | Pedunculopontine nucleus | Brainstem+Pons |
| Schneider et al 2021 | Medial Leminscus WM) | Brainstem+Pons |
| Inglese et al 2010 | WM, Frontal | EXCLUDED |
| Inglese et al 2010 | WM, Occipital | EXCLUDED |
| Reetz et al 2012 | Insula | EXCLUDED |
| Reetz et al 2012 | Cingulate | EXCLUDED |
| Reetz et al 2012 | GM, Cortex, Pre-/Postcentral | EXCLUDED |
| Paling et al 2013 | DeepGM caudate, putamen, pallidum, thalamus) | EXCLUDED |
| Mirkes et al 2015 | WM, prefrontal | EXCLUDED |
| Thulborn et al 2016 | Basal Ganglia | EXCLUDED |
| Liao et al 2019 | Insula | EXCLUDED |
| Gerhalter et al 2021 | WM, Frontal | EXCLUDED |
| Gerhalter et al 2021 | WM, posterior | EXCLUDED |
| Brownlee et al 2019 | Deep grey matter | EXCLUDED |
| Schneider et al 2021 | Nucleus Accumbens | EXCLUDED |
| Schneider et al 2021 | Bed Nucleus of stria terminalis, Fibres, Per-ventricular surface of the thalamus | EXCLUDED |
| Schneider et al 2021 | Substantia nigra | EXCLUDED |
| Schneider et al 2021 | Subthalamic Nucleus | EXCLUDED |
| Schneider et al 2021 | Cerebral peduncle WM) | EXCLUDED |
| Schneider et al 2021 | Mammillary Body | EXCLUDED |
| Schneider et al 2021 | Habenula epithalamic / just above thal) | EXCLUDED |
| Schneider et al 2021 | Nucleus basalis Meynert | EXCLUDED |

#### Supplementary Table 6 – Effect of moderators in intraregional variability (Hedges' g)

| | | Sequence | Comparison group | Calibration method | Voxel volume | Field strength (Tesla) | Repetition time (TR) | Echo time |
|---|---|---|---|---|---|---|---|---|
| GM | SMD | -2.58 | -1.62 | -0.63 | -0.03 | 0.28 | 0.02 | -0.35 |
| | -95% CI | -3.37 | -2.29 | -1.22 | -0.62 | -0.3 | -0.56 | -0.92 |
| | +95% CI | -1.76 | -0.93 | -0.04 | 0.54 | 0.87 | 0.6 | -0.24 |
| Temporal GM | SMD | -3.67 | 0.43 | 3.39 | 0.01 | -0.04 | -2.52 | -5.24 |
| | -95% CI | -5.19 | -0.42 | 1.96 | -0.74 | -0.88 | -3.68 | -7.27 |
| | +95% CI | -2.12 | 1.28 | 4.78 | 0.94 | 0.8 | -1.31 | -3.19 |
| Thalamus | SMD | 6.54 | 5.82 | 6.78 | 2.62 | 0.6 | -3.66 | 0.22 |
| | -95% CI | 4.04 | 3.59 | 4.2 | 1.39 | -0.27 | -5.21 | 0.62 |
| | +95% CI | 9.01 | 8.02 | 9.34 | 3.81 | 1.45 | -2.07 | 1.06 |
| Brainstem +Pons | SMD | 1.22 | -1.23 | 0.94 | -1.23 | -0.31 | 0.39 | 0.29 |
| | -95% CI | 0.42 | -2.01 | 0.17 | -2.02 | -1.03 | -0.34 | -0.43 |
| | +95% CI | 2.01 | -0.42 | 1.7 | -0.43 | 0.41 | 1.11 | 1.01 |
| WM | SMD | -4.28 | -3.42 | 0.61 | -0.43 | 0.62 | -0.28 | 1.75 |
| | -95% CI | -5.38 | -4.36 | 0.01 | -1.02 | 0.02 | -0.87 | 1.05 |
| | +95% CI | -3.17 | -2.4 | 1.2 | 0.16 | 1.21 | 0.3 | 2.43 |
| 'Central' WM | SMD | 2.79 | 2.99 | 1.19 | 0 | -0.23 | -0.72 | -4.17 |
| | -95% CI | 1.53 | 1.68 | 0.25 | -0.84 | -1.07 | -1.58 | -5.77 |
| | +95% CI | 4.02 | 4.26 | 2.1 | 0.84 | 0.62 | 0.16 | -2.53 |

#### Supplementary Table 7 – Data Table

BT, Brain Tumour; CSF, Cerebrospinal Fluid (Ventricular); Ext, External; HD, Huntingdon's Disease ; M, Migraine; MS, Multiple Sclerosis; S, Stroke; TBI, Traumatic Brain Injury; VH, Vitreous Humour;

* except for Zaaraoui et 2012 who reported range and Driver et al 2019 who reported standard error

| Paper | Tesla | Sequence | Calibration | TR (ms) | TE (ms) | X (mm) | Y (mm) | Z (mm) | Voxel volume (mm3) | N | Mean Age (years) | S.D. (Age) | Sex (N Female) | Comparison group | Tissue | Mean TSC (mM) | TSC S.D* |
|---|---|---|---|---|---|---|---|---|---|---|---|---|---|---|---|---|---|
| Winkler et al 1989 | 1.5 | GRE | VH | 98 | 3 | 7 | 7 | 10 | 490 | 4 | NR | NR | NR | No | Deep WM | 34 | 7 |
| Winkler et al 1989 | 1.5 | GRE | VH | 98 | 3 | 7 | 7 | 10 | 490 | 4 | NR | NR | NR | No | GM | 69 | 16 |
| Ouwerkerk et al 2003 | 1.5 | TPI | Ext | 120 | 0.37 | 3.4 | 3.4 | 3.4 | 39.304 | 9 | NR | NR | 3 | BT | GM | 60 | 6 |
| Ouwerkerk et al 2003 | 1.5 | TPI | Ext | 120 | 0.37 | 3.4 | 3.4 | 3.4 | 39.304 | 9 | NR | NR | 3 | BT | WM | 69 | 11 |
| Thulborn et al 2005 | 3 | TPI | Ext | 100 | 0.3 | 5 | 5 | 5 | 125 | 5 | NR | NR | NR | S | GM. Frontal | 37.5 | 2.8 |
| Thulborn et al 2005 | 3 | TPI | Ext | 100 | 0.3 | 5 | 5 | 5 | 125 | 5 | NR | NR | NR | S | GM. Frontal | 37.9 | 2.6 |
| Thulborn et al 2005 | 3 | TPI | Ext | 100 | 0.3 | 5 | 5 | 5 | 125 | 5 | NR | NR | NR | S | GM. Parietal | 38.1 | 3.1 |
| Thulborn et al 2005 | 3 | TPI | Ext | 100 | 0.3 | 5 | 5 | 5 | 125 | 5 | NR | NR | NR | S | GM. Parietal | 42.2 | 2.5 |
| Thulborn et al 2005 | 3 | TPI | Ext | 100 | 0.3 | 5 | 5 | 5 | 125 | 5 | NR | NR | NR | S | GM. Temporal | 42.1 | 3 |
| Thulborn et al 2005 | 3 | TPI | Ext | 100 | 0.3 | 5 | 5 | 5 | 125 | 5 | NR | NR | NR | S | GM. Temporal | 41.4 | 2.8 |
| Thulborn et al 2005 | 3 | TPI | Ext | 100 | 0.3 | 5 | 5 | 5 | 125 | 5 | NR | NR | NR | S | GM. Occipital | 41.7 | 1.6 |
| Thulborn et al 2005 | 3 | TPI | Ext | 100 | 0.3 | 5 | 5 | 5 | 125 | 5 | NR | NR | NR | S | GM. Occipital | 41.8 | 1.1 |
| Thulborn et al 2005 | 3 | TPI | Ext | 100 | 0.3 | 5 | 5 | 5 | 125 | 5 | NR | NR | NR | S | GM | 39.9 | 2.4 |
| Thulborn et al 2005 | 3 | TPI | Ext | 100 | 0.3 | 5 | 5 | 5 | 125 | 5 | NR | NR | NR | S | GM | 40.8 | 2 |
| Thulborn et al 2005 | 3 | TPI | Ext | 100 | 0.3 | 5 | 5 | 5 | 125 | 5 | NR | NR | NR | S | Central WM | 35.5 | 2.1 |
| Thulborn et al 2005 | 3 | TPI | Ext | 100 | 0.3 | 5 | 5 | 5 | 125 | 5 | NR | NR | NR | S | Central WM | 37 | 1.3 |
| Thulborn et al 2005 | 3 | TPI | Ext | 100 | 0.3 | 5 | 5 | 5 | 125 | 5 | NR | NR | NR | S | Central WM | 34.2 | 3 |
| Thulborn et al 2005 | 3 | TPI | Ext | 100 | 0.3 | 5 | 5 | 5 | 125 | 5 | NR | NR | NR | S | Central WM | 35.9 | 2.5 |
| Thulborn et al 2005 | 3 | TPI | Ext | 100 | 0.3 | 5 | 5 | 5 | 125 | 5 | NR | NR | NR | S | Deep WM | 33.2 | 3.1 |
| Thulborn et al 2005 | 3 | TPI | Ext | 100 | 0.3 | 5 | 5 | 5 | 125 | 5 | NR | NR | NR | S | Deep WM | 33.2 | 3.1 |

| | | | | | | | | | | | | | | | | | |
|---|---|---|---|---|---|---|---|---|---|---|---|---|---|---|---|---|---|
| Thulborn et al 2005 | 3 | TPI | Ext | 100 | 0.3 | 5 | 5 | 5 | 125 | 5 | NR | NR | NR | S | Deep WM | 28.1 | 3.4 |
| Thulborn et al 2005 | 3 | TPI | Ext | 100 | 0.3 | 5 | 5 | 5 | 125 | 5 | NR | NR | NR | S | Deep WM | 27.8 | 3.5 |
| Thulborn et al 2005 | 3 | TPI | Ext | 100 | 0.3 | 5 | 5 | 5 | 125 | 5 | NR | NR | NR | S | WM | 32.8 | 3.2 |
| Thulborn et al 2005 | 3 | TPI | Ext | 100 | 0.3 | 5 | 5 | 5 | 125 | 5 | NR | NR | NR | S | WM | 33.5 | 4.1 |
| Thulborn et al 2005 | 3 | TPI | Ext | 100 | 0.3 | 5 | 5 | 5 | 125 | 5 | NR | NR | NR | S | Thalamus | 32.8 | 3.2 |
| Thulborn et al 2005 | 3 | TPI | Ext | 100 | 0.3 | 5 | 5 | 5 | 125 | 5 | NR | NR | NR | S | Thalamus | 32.1 | 2.6 |
| Thulborn et al 2005 | 3 | TPI | Ext | 100 | 0.3 | 5 | 5 | 5 | 125 | 5 | NR | NR | NR | S | Putamen | 37.3 | 4.1 |
| Thulborn et al 2005 | 3 | TPI | Ext | 100 | 0.3 | 5 | 5 | 5 | 125 | 5 | NR | NR | NR | S | Putamen | 39.1 | 3.2 |
| Thulborn et al 2005 | 3 | TPI | Ext | 100 | 0.3 | 5 | 5 | 5 | 125 | 5 | NR | NR | NR | S | WM. Cb+DN | 35.5 | 2.6 |
| Thulborn et al 2005 | 3 | TPI | Ext | 100 | 0.3 | 5 | 5 | 5 | 125 | 5 | NR | NR | NR | S | WM. Cb+DN | 33.6 | 2.1 |
| Thulborn et al 2005 | 3 | TPI | Ext | 100 | 0.3 | 5 | 5 | 5 | 125 | 5 | NR | NR | NR | S | GM. Temporal | 47.1 | 2.5 |
| Thulborn et al 2005 | 3 | TPI | Ext | 100 | 0.3 | 5 | 5 | 5 | 125 | 5 | NR | NR | NR | S | GM. Temporal | 46.7 | 2.1 |
| Thulborn et al 2005 | 3 | TPI | Ext | 100 | 0.3 | 5 | 5 | 5 | 125 | 5 | NR | NR | NR | S | Brainstem +Pons | 36.7 | 1.6 |
| Thulborn et al 2005 | 3 | TPI | Ext | 100 | 0.3 | 5 | 5 | 5 | 125 | 5 | NR | NR | NR | S | Brainstem +Pons | 37 | 1.9 |
| Thulborn et al 2005 | 3 | TPI | Ext | 100 | 0.3 | 5 | 5 | 5 | 125 | 5 | NR | NR | NR | S | Brainstem +Pons | 31.3 | 1.5 |
| Thulborn et al 2005 | 3 | TPI | Ext | 100 | 0.3 | 5 | 5 | 5 | 125 | 5 | NR | NR | NR | S | Brainstem +Pons | 30.9 | 2.9 |
| Inglese et al 2010 | 3 | Radial | Ext | 120 | 0.05 | 4 | 4 | 4 | 64 | 13 | 36.7 | NR | 10 | MS | Central WM | 20.26 | 2.35 |
| Inglese et al 2010 | 3 | Radial | Ext | 120 | 0.05 | 4 | 4 | 4 | 64 | 13 | 36.7 | NR | 10 | MS | Central WM | 17.22 | 2.7 |
| Inglese et al 2010 | 3 | Radial | Ext | 120 | 0.05 | 4 | 4 | 4 | 64 | 13 | 36.7 | NR | 10 | MS | WM | 19.38 | 1.65 |
| Inglese et al 2010 | 3 | Radial | Ext | 120 | 0.05 | 4 | 4 | 4 | 64 | 13 | 36.7 | NR | 10 | MS | WM. Cb+DN | 21.28 | 3.62 |
| Inglese et al 2010 | 3 | Radial | Ext | 120 | 0.05 | 4 | 4 | 4 | 64 | 13 | 36.7 | NR | 10 | MS | GM. Frontal | 30.22 | 4.77 |
| Inglese et al 2010 | 3 | Radial | Ext | 120 | 0.05 | 4 | 4 | 4 | 64 | 13 | 36.7 | NR | 10 | MS | GM. Occipital | 30.34 | 6.28 |
| Inglese et al 2010 | 3 | Radial | Ext | 120 | 0.05 | 4 | 4 | 4 | 64 | 13 | 36.7 | NR | 10 | MS | GM. Parietal | 30 | 4.14 |
| Inglese et al 2010 | 3 | Radial | Ext | 120 | 0.05 | 4 | 4 | 4 | 64 | 13 | 36.7 | NR | 10 | MS | GM. Temporal | 31.58 | 4.3 |
| Inglese et al 2010 | 3 | Radial | Ext | 120 | 0.05 | 4 | 4 | 4 | 64 | 13 | 36.7 | NR | 10 | MS | GM | 30.54 | 3.17 |
| Lu et al 2010 | 3 | FlexTPI | Ext | 160 | 0.36 | 5 | 5 | 5 | 125 | 5 | 32.4 | 8.9 | 1 | No | GM | 38.1 | 0.6 |
| Lu et al 2010 | 3 | FlexTPI | Ext | 160 | 0.36 | 5 | 5 | 5 | 125 | 5 | 32.4 | 8.9 | 1 | No | WM | 28.7 | 1.2 |

| | | | | | | | | | | | | | | | | | |
|---|---|---|---|---|---|---|---|---|---|---|---|---|---|---|---|---|---|
| Reetz et al 2012 | 4 | SPRITE | Ext | 10 | 0.3 | 4 | 4 | 4 | 64 | 13 | 44.9 | 9.9 | 6 | HD | Caudate | 52 | 6 |
| Reetz et al 2012 | 4 | SPRITE | Ext | 10 | 0.3 | 4 | 4 | 4 | 64 | 13 | 44.9 | 9.9 | 6 | HD | Putamen | 46 | 3 |
| Reetz et al 2012 | 4 | SPRITE | Ext | 10 | 0.3 | 4 | 4 | 4 | 64 | 13 | 44.9 | 9.9 | 6 | HD | Globus Pallidus | 44 | 4 |
| Reetz et al 2012 | 4 | SPRITE | Ext | 10 | 0.3 | 4 | 4 | 4 | 64 | 13 | 44.9 | 9.9 | 6 | HD | Thalamus | 49 | 4 |
| Reetz et al 2012 | 4 | SPRITE | Ext | 10 | 0.3 | 4 | 4 | 4 | 64 | 13 | 44.9 | 9.9 | 6 | HD | GM. Temporal | 50 | 5 |
| Reetz et al 2012 | 4 | SPRITE | Ext | 10 | 0.3 | 4 | 4 | 4 | 64 | 13 | 44.9 | 9.9 | 6 | HD | GM. Temporal | 55 | 5 |
| Reetz et al 2012 | 4 | SPRITE | Ext | 10 | 0.3 | 4 | 4 | 4 | 64 | 13 | 44.9 | 9.9 | 6 | HD | GM. Parietal | 55 | 5 |
| Reetz et al 2012 | 4 | SPRITE | Ext | 10 | 0.3 | 4 | 4 | 4 | 64 | 13 | 44.9 | 9.9 | 6 | HD | GM. Frontal | 48 | 5 |
| Reetz et al 2012 | 4 | SPRITE | Ext | 10 | 0.3 | 4 | 4 | 4 | 64 | 13 | 44.9 | 9.9 | 6 | HD | GM. Temporal | 50 | 4 |
| Reetz et al 2012 | 4 | SPRITE | Ext | 10 | 0.3 | 4 | 4 | 4 | 64 | 13 | 44.9 | 9.9 | 6 | HD | GM. Parietal | 49 | 6 |
| Reetz et al 2012 | 4 | SPRITE | Ext | 10 | 0.3 | 4 | 4 | 4 | 64 | 13 | 44.9 | 9.9 | 6 | HD | GM. Occipital | 51 | 3 |
| Reetz et al 2012 | 4 | SPRITE | Ext | 10 | 0.3 | 4 | 4 | 4 | 64 | 13 | 44.9 | 9.9 | 6 | HD | WM. Cb+DN | 38 | 3 |
| Qian et al 2012 | 7 | AWSOS | CSF | 100 | 0.5 | 0.86 | 0.86 | 4 | 2.9584 | 5 | NR | NR | 5 | No | GM | 47.1 | 6.7 |
| Qian et al 2012 | 7 | AWSOS | CSF | 100 | 0.5 | 0.86 | 0.86 | 4 | 2.9584 | 5 | NR | NR | 5 | No | WM | 37.4 | 2.9 |
| Zaaraoui et 2012 | 3 | DA Radial | Ext | 120 | 0.2 | 3.6 | 3.6 | 3.6 | 46.656 | 15 | NR | NR | 12 | MS | GM | 53.5 | 8.55 |
| Zaaraoui et 2012 | 3 | DA Radial | Ext | 120 | 0.2 | 3.6 | 3.6 | 3.6 | 46.656 | 15 | NR | NR | 12 | MS | WM | 48.8 | 7.65 |
| Paling et al 2013 | 3 | Radial | Ext | 120 | 0.27 | 4 | 4 | 4 | 64 | 27 | 42.9 | 11.3 | 16 | MS | GM | 36 | 2.7 |
| Paling et al 2013 | 3 | Radial | Ext | 120 | 0.27 | 4 | 4 | 4 | 64 | 27 | 42.9 | 11.3 | 16 | MS | WM | 32.1 | 2.2 |
| Maarouf et al 2014 | 3 | DA Radial | Ext | 120 | 0.2 | 3.6 | 3.6 | 3.6 | 46.656 | 15 | NR | NR | NR | MS | GM | 48.1 | 2.3 |
| Maarouf et al 2014 | 3 | DA Radial | Ext | 120 | 0.2 | 3.6 | 3.6 | 3.6 | 46.656 | 15 | NR | NR | NR | MS | WM | 41.7 | 2.2 |
| Mirkes et al 2015 | 9.4 | AWSOS | Ext | 150 | 0.3 | 1 | 1 | 5 | 5 | 5 | 29 | 4 | 1 | No | GM. Frontal | 36 | 2 |
| Niesporak et al 2015 | 7 | DA Radial | Ext | 150 | 0.45 | 3 | 3 | 3 | 27 | 4 | 26 | 2 | 1 | No | GM | 48 | 1 |
| Niesporak et al 2015 | 7 | DA Radial | Ext | 150 | 0.45 | 3 | 3 | 3 | 27 | 4 | 26 | 2 | 1 | No | WM | 41 | 3 |
| Eisele et al 2016 | 3 | DA Radial | Ext | 60 | 0.22 | 3.6 | 3.6 | 3.6 | 46.656 | 10 | NR | NR | 5 | MS | GM | 40.09 | 4.64 |
| Eisele et al 2016 | 3 | DA Radial | Ext | 60 | 0.22 | 3.6 | 3.6 | 3.6 | 46.656 | 10 | NR | NR | 5 | MS | WM | 35.17 | 3.4 |
| Petracca et al 2016 | 7 | GRE | Ext | 150 | 6.8 | 5 | 5 | 5 | 125 | 17 | 46.16 | 11.65 | 8 | MS | GM | 40.26 | 2.99 |
| Petracca et al 2016 | 7 | GRE | Ext | 150 | 6.8 | 5 | 5 | 5 | 125 | 17 | 46.16 | 11.65 | 8 | MS | WM | 27.88 | 2.55 |

| | | | | | | | | | | | | | | | | | |
|---|---|---|---|---|---|---|---|---|---|---|---|---|---|---|---|---|---|
| Thulborn et al 2016 | 9.4 | FlexTPI | Ext | 160 | 0.26 | 3.5 | 3.5 | 3.5 | 42.875 | 45 | 48 | 19 | NR | No | Brainstem +Pons | 36.3 | 1.5 |
| Thulborn et al 2016 | 9.4 | FlexTPI | Ext | 160 | 0.26 | 3.5 | 3.5 | 3.5 | 42.875 | 45 | 48 | 19 | NR | No | Thalamus | 37.1 | 2.4 |
| Maarouf et al 2017 | 3 | DA Radial | Ext | 120 | 0.2 | 3.6 | 3.6 | 3.6 | 46.656 | 31 | 32.7 | 12.4 | 15 | MS | GM | 52.7 | 3.6 |
| Maarouf et al 2017 | 3 | DA Radial | Ext | 120 | 0.2 | 3.6 | 3.6 | 3.6 | 46.656 | 31 | 35.7 | 12.4 | 15 | MS | WM | 45.9 | 3.1 |
| Eisele et al 2017 | 3 | DA Radial | Ext | 60 | 0.22 | 3.6 | 3.6 | 3.6 | 46.656 | 6 | 42 | 10 | 5 | MS | WM. Cb+DN | 33.14 | 1.12 |
| Eisele et al 2017 | 3 | DA Radial | Ext | 60 | 0.22 | 3.6 | 3.6 | 3.6 | 46.656 | 6 | 42 | 10 | 5 | MS | Globus Pallidus | 33.24 | 2.29 |
| Ridley et al 2018 | 7 | DA Radial | Ext | 120 | 0.3 | 3.5 | 3.5 | 3.5 | 42.875 | 13 | 23.9 | 3.6 | 5 | No | Thalamus | 40.3 | 2.33 |
| Ridley et al 2018 | 7 | DA Radial | Ext | 120 | 0.3 | 3.5 | 3.5 | 3.5 | 42.875 | 13 | 23.9 | 3.6 | 5 | No | Putamen | 36.29 | 1.76 |
| Ridley et al 2018 | 7 | DA Radial | Ext | 120 | 0.3 | 3.5 | 3.5 | 3.5 | 42.875 | 13 | 23.9 | 3.6 | 5 | No | Globus Pallidus | 33.24 | 2.62 |
| Ridley et al 2018 | 7 | DA Radial | Ext | 120 | 0.3 | 3.5 | 3.5 | 3.5 | 42.875 | 13 | 23.9 | 3.6 | 5 | No | Caudate | 37.06 | 2.09 |
| Ridley et al 2018 | 7 | DA Radial | Ext | 120 | 0.3 | 3.5 | 3.5 | 3.5 | 42.875 | 13 | 23.9 | 3.6 | 5 | No | GM | 46.92 | 2.83 |
| Ridley et al 2018 | 7 | DA Radial | Ext | 120 | 0.3 | 3.5 | 3.5 | 3.5 | 42.875 | 13 | 23.9 | 3.6 | 5 | No | Central WM | 35.52 | 3.35 |
| Ridley et al 2018 | 7 | DA Radial | Ext | 120 | 0.3 | 3.5 | 3.5 | 3.5 | 42.875 | 13 | 23.9 | 3.6 | 5 | No | WM. Cb+DN | 28.6 | 2.88 |
| Ridley et al 2018 | 7 | DA Radial | Ext | 120 | 0.3 | 3.5 | 3.5 | 3.5 | 42.875 | 13 | 23.9 | 3.6 | 5 | No | Deep WM | 32.39 | 2.74 |
| Ridley et al 2018 | 7 | DA Radial | Ext | 120 | 0.3 | 3.5 | 3.5 | 3.5 | 42.875 | 13 | 23.9 | 3.6 | 5 | No | Brainstem +Pons | 25.18 | 3.17 |
| Ridley et al 2018 | 7 | DA Radial | Ext | 120 | 0.3 | 3.5 | 3.5 | 3.5 | 42.875 | 13 | 23.9 | 3.6 | 5 | No | WM | 38.15 | 1.97 |
| Worthoff et al 2018 | 4 | SISTINA | VH | 150 | 0.36 | 6 | 6 | 6 | 216 | 40 | NR | NR | 16 | No | WM | 25 | 1 |
| Riemer et al 2019 | 3 | Cones | Ext | 100 | 0.5 | 4 | 4 | 4 | 64 | 11 | 32 | 6 | 3 | No | GM | 52.1 | 7.1 |
| Riemer et al 2019 | 3 | Cones | Ext | 100 | 0.5 | 4 | 4 | 4 | 64 | 11 | 32 | 6 | 3 | No | WM | 41.8 | 6.7 |
| Driver et al 2019 | 4.7 | TPI | VH | 85 | 0.11 | 3.2 | 3.2 | 6.4 | 65.536 | 9 | 30 | 6 | 5 | No | GM | 41.9 | 0.9 |
| Driver et al 2019 | 4.7 | TPI | VH | 85 | 0.11 | 3.2 | 3.2 | 6.4 | 65.536 | 9 | 30 | 6 | 5 | No | WM | 39.1 | 0.8 |
| Liao et al 2019 | 3 | TPI | CSF | 160 | 0.4 | 3.44 | 3.44 | 3.44 | 40.70758 | 8 | NR | NR | 3 | No | GM | 45 | 8 |
| Liao et al 2019 | 3 | TPI | CSF | 160 | 0.4 | 3.44 | 3.44 | 3.44 | 40.70758 | 8 | NR | NR | 3 | No | GM. Frontal | 48 | 6 |
| Liao et al 2019 | 3 | TPI | CSF | 160 | 0.4 | 3.44 | 3.44 | 3.44 | 40.70758 | 8 | NR | NR | 3 | No | GM. Occipital | 48 | 6 |
| Liao et al 2019 | 3 | TPI | CSF | 160 | 0.4 | 3.44 | 3.44 | 3.44 | 40.70758 | 8 | NR | NR | 3 | No | GM. Parietal | 47 | 6 |
| Liao et al 2019 | 3 | TPI | CSF | 160 | 0.4 | 3.44 | 3.44 | 3.44 | 40.70758 | 8 | NR | NR | 3 | No | GM. Temporal | 47 | 6 |
| Liao et al 2019 | 3 | TPI | CSF | 160 | 0.4 | 3.44 | 3.44 | 3.44 | 40.70758 | 8 | NR | NR | 3 | No | GM. Temporal | 44 | 7 |

| | | | | | | | | | | | | | | | | | |
|---|---|---|---|---|---|---|---|---|---|---|---|---|---|---|---|---|---|
| Liao et al 2019 | 3 | TPI | CSF | 160 | 0.4 | 3.44 | 3.44 | 3.44 | 40.70758 | 8 | NR | NR | 3 | No | Thalamus | 38 | 4 |
| Liao et al 2019 | 3 | TPI | CSF | 160 | 0.4 | 3.44 | 3.44 | 3.44 | 40.70758 | 8 | NR | NR | 3 | No | Putamen | 42 | 7 |
| Liao et al 2019 | 3 | TPI | CSF | 160 | 0.4 | 3.44 | 3.44 | 3.44 | 40.70758 | 8 | NR | NR | 3 | No | Caudate | 43 | 4 |
| Liao et al 2019 | 3 | TPI | CSF | 160 | 0.4 | 3.44 | 3.44 | 3.44 | 40.70758 | 8 | NR | NR | 3 | No | WM | 41 | 7 |
| Meyer et al 2019a | 3 | DA Radial | Ext | 120 | 0.2 | 3.6 | 3.6 | 3.6 | 46.656 | 12 | 31 | 8.3 | 8 | No | GM | 51.5 | 4.5 |
| Meyer et al 2019a | 3 | DA Radial | Ext | 120 | 0.2 | 3.6 | 3.6 | 3.6 | 46.656 | 12 | 31 | 8.3 | 8 | No | WM | 40.9 | 3.8 |
| Meyer et al 2019a | 3 | DA Radial | Ext | 120 | 0.2 | 3.6 | 3.6 | 3.6 | 46.656 | 12 | 31 | 8.3 | 8 | No | Caudate | 60.9 | 8.1 |
| Meyer et al 2019a | 3 | DA Radial | Ext | 120 | 0.2 | 3.6 | 3.6 | 3.6 | 46.656 | 12 | 31 | 8.3 | 8 | No | Brainstem +Pons | 39.8 | 5.3 |
| Meyer et al 2019a | 3 | DA Radial | Ext | 120 | 0.2 | 3.6 | 3.6 | 3.6 | 46.656 | 12 | 31 | 8.3 | 8 | No | WM. Cb+DN | 40.1 | 4.9 |
| Meyer et al 2019b | 3 | DA Radial | Ext | 120 | 0.2 | 4 | 4 | 4 | 64 | 12 | 34.3 | 10.7 | 12 | M | GM. Parietal | 39.9 | 1.6 |
| Meyer et al 2019b | 3 | DA Radial | Ext | 120 | 0.2 | 4 | 4 | 4 | 64 | 12 | 34.3 | 10.7 | 12 | M | Deep WM | 32.8 | 2.2 |
| Meyer et al 2019b | 3 | DA Radial | Ext | 120 | 0.2 | 4 | 4 | 4 | 64 | 12 | 34.3 | 10.7 | 12 | M | WM. Cb+DN | 32.8 | 2.3 |
| Meyer et al 2019b | 3 | DA Radial | Ext | 120 | 0.2 | 4 | 4 | 4 | 64 | 12 | 34.3 | 10.7 | 12 | M | Brainstem +Pons | 30.4 | 2.5 |
| Kim et al 2020 | 7 | GRE | CSF | 100 | 4 | 4 | 4 | 4 | 64 | 8 | NR | NR | 0 | No | GM | 58.72 | 4.39 |
| Kim et al 2020 | 7 | GRE | CSF | 100 | 4 | 4 | 4 | 4 | 64 | 8 | NR | NR | 0 | No | WM | 35.61 | 2.82 |
| Gerhalter et al 2021 | 3 | FLORET | VH | 100 | 0.2 | 6 | 6 | 6 | 216 | 19 | 31.4 | 7.5 | 12 | TBI | WM | 38.4 | 3 |
| Gerhalter et al 2021 | 3 | FLORET | VH | 100 | 0.2 | 6 | 6 | 6 | 216 | 19 | 31.4 | 7.5 | 12 | TBI | Central WM | 45.7 | 5.6 |
| Gerhalter et al 2021 | 3 | FLORET | VH | 100 | 0.2 | 6 | 6 | 6 | 216 | 19 | 31.4 | 7.5 | 12 | TBI | Central WM | 46.7 | 4 |
| Gerhalter et al 2021 | 3 | FLORET | VH | 100 | 0.2 | 6 | 6 | 6 | 216 | 19 | 31.4 | 7.5 | 12 | TBI | Central WM | 38.9 | 4.8 |
| Gerhalter et al 2021 | 3 | FLORET | VH | 100 | 0.2 | 6 | 6 | 6 | 216 | 19 | 31.4 | 7.5 | 12 | TBI | Deep WM | 30 | 2 |
| Gerhalter et al 2021 | 3 | FLORET | VH | 100 | 0.2 | 6 | 6 | 6 | 216 | 19 | 31.4 | 7.5 | 12 | TBI | GM | 43.2 | 3.8 |
| Gerhalter et al 2021 | 3 | FLORET | VH | 100 | 0.2 | 6 | 6 | 6 | 216 | 19 | 31.4 | 7.5 | 12 | TBI | Caudate | 48.5 | 3.5 |
| Gerhalter et al 2021 | 3 | FLORET | VH | 100 | 0.2 | 6 | 6 | 6 | 216 | 19 | 31.4 | 7.5 | 12 | TBI | Globus Pallidus | 30.2 | 2.2 |
| Gerhalter et al 2021 | 3 | FLORET | VH | 100 | 0.2 | 6 | 6 | 6 | 216 | 19 | 31.4 | 7.5 | 12 | TBI | Putamen | 33.2 | 2.3 |
| Gerhalter et al 2021 | 3 | FLORET | VH | 100 | 0.2 | 6 | 6 | 6 | 216 | 19 | 31.4 | 7.5 | 12 | TBI | Thalamus | 40.2 | 4 |
| Brownlee et al 2019 | 3 | Cones | Ext | 120 | 0.22 | 3 | 3 | 3 | 27 | 34 | 35.5 | 10.1 | 23 | MS | GM | 40.63 | 2.33 |
| Brownlee et al 2019 | 3 | Cones | Ext | 120 | 0.22 | 3 | 3 | 3 | 27 | 34 | 35.5 | 10.1 | 23 | MS | WM | 32.03 | 3.44 |

| | | | | | | | | | | | | | | | | | |
|---|---|---|---|---|---|---|---|---|---|---|---|---|---|---|---|---|---|
| Schneider et al 2021 | 7 | DA Radial | CSF | 100 | 0.35 | 2 | 2 | 2 | 8 | 5 | 28.4 | 6.5 | 3 | No | Caudate | 53.1 | 4.2 |
| Schneider et al 2021 | 7 | DA Radial | CSF | 100 | 0.35 | 2 | 2 | 2 | 8 | 5 | 28.4 | 6.5 | 3 | No | Brainstem +Pons | 38.9 | 1.2 |
| Schneider et al 2021 | 7 | DA Radial | CSF | 100 | 0.35 | 2 | 2 | 2 | 8 | 5 | 28.4 | 6.5 | 3 | No | Globus Pallidus | 38.8 | 2 |
| Schneider et al 2021 | 7 | DA Radial | CSF | 100 | 0.35 | 2 | 2 | 2 | 8 | 5 | 28.4 | 6.5 | 3 | No | Putamen | 47.1 | 2.9 |
| Schneider et al 2021 | 7 | DA Radial | CSF | 100 | 0.35 | 2 | 2 | 2 | 8 | 5 | 28.4 | 6.5 | 3 | No | Brainstem +Pons | 80.3 | 7.3 |
| Schneider et al 2021 | 7 | DA Radial | CSF | 100 | 0.35 | 2 | 2 | 2 | 8 | 5 | 28.4 | 6.5 | 3 | No | Brainstem +Pons | 43 | 2.8 |
| Schneider et al 2021 | 7 | DA Radial | CSF | 100 | 0.35 | 2 | 2 | 2 | 8 | 5 | 28.4 | 6.5 | 3 | No | Brainstem +Pons | 53.7 | 3.6 |
| Schneider et al 2021 | 7 | DA Radial | CSF | 100 | 0.35 | 2 | 2 | 2 | 8 | 5 | 28.4 | 6.5 | 3 | No | Thalamus | 45.8 | 3.7 |
| Schneider et al 2021 | 7 | DA Radial | CSF | 100 | 0.35 | 2 | 2 | 2 | 8 | 5 | 28.4 | 6.5 | 3 | No | Thalamus | 57.2 | 5.5 |
| Schneider et al 2021 | 7 | DA Radial | CSF | 100 | 0.35 | 2 | 2 | 2 | 8 | 5 | 28.4 | 6.5 | 3 | No | Brainstem +Pons | 46.4 | 4.5 |
| Schneider et al 2021 | 7 | DA Radial | CSF | 100 | 0.35 | 2 | 2 | 2 | 8 | 5 | 28.4 | 6.5 | 3 | No | Thalamus | 49.8 | 4.7 |
| Schneider et al 2021 | 7 | DA Radial | CSF | 100 | 0.35 | 2 | 2 | 2 | 8 | 5 | 28.4 | 6.5 | 3 | No | Globus Pallidus | 40.9 | 2.2 |
| Schneider et al 2021 | 7 | DA Radial | CSF | 100 | 0.35 | 2 | 2 | 2 | 8 | 5 | 28.4 | 6.5 | 3 | No | Brainstem +Pons | 46.4 | 3.4 |
